# No evidence of phago-mixotropy in *Micromonas polaris*, the dominant picophytoplankton species in the Arctic

**DOI:** 10.1101/2020.05.26.117895

**Authors:** Valeria Jimenez, John A. Burns, Florence Le Gall, Fabrice Not, Daniel Vaulot

**Affiliations:** Sorbonne Université, CNRS, UMR7144, Ecology of Marine Plankton team, Station Biologique de Roscoff, 29680 Roscoff, France; Bigelow Laboratory for Ocean Sciences, East Boothbay, ME, USA; Asian School of the Environment, Nanyang Technological University, 50 Nanyang Avenue, Singapore 639798

**Keywords:** phytoplankton, Arctic, phago-mixotrophy, *Micromonas*

## Abstract

In the Arctic Ocean, the small green alga *Micromonas polaris* dominates pico-phytoplankton during the summer months. It has been previously hypothesized to be phago-mixotrophic (capable of bacteria ingestion) based on laboratory and field experiments. Prey uptake was analysed in several *M. polaris* strains isolated from different regions and depths of the Arctic Ocean. Using both fluorescent beads and fluorescently labelled bacteria as prey, we found no evidence of phago-mixotrophy in any *M. polaris* strain by flow cytometric measurement of prey ingestion. In addition, *in silico* predictions reveal that members of the genus *Micromonas* lack a genetic signature of phagocytotic capacity.

## Introduction

Polar regions are undergoing drastic changes due to climate change and global warming in particular. These changes have strong effects in Arctic marine ecosystems [1–3] where phytoplankton production plays an essential role in food web dynamics and biogeochemical cycles [4–6]. Considerable spatial and temporal changes in primary production have been observed in the last two decades [5, 7, 8]. Rapid melting and early ice retreat increase the open areas exposed to solar radiation which in turns results in a considerable increase in annual net primary production along with a lengthening of the phytoplankton growing season [5, 6, 8]. Changes in Arctic primary production are also influenced by the increase of freshwater delivery to the upper ocean that leads to stronger water column stratification limiting the upward flux of nutrients to the surface [6, 9–14].

Our ability to explain and predict the responses of Arctic phytoplankton communities to climate change is challenged by our limited understanding regarding their ecological and physiological strategies of growth and survival. Arctic phytoplankton communities experience extreme environmental conditions such as nutrient limitation, exposure to a long period of darkness (polar winter) and changes in light levels under the ice caused by the variation in snow coverage and ice thickness [15]. In such a highly variable context, it has been suggested that phago-mixotrophy (ability to combine photosynthesis and bacterivory) could be a common trophic strategy among Arctic protists [16]. At the scale of the global ocean, phago-mixotrophy is an important, but until recently underestimated, process for energy and nutrient transfer (e.g. carbon fluxes) throughout the food web [17–19]. Phago-mixotrophic plankton are widespread in the ocean and evolutionary diverse, spread across the tree of life [20]. They account for a large proportion of bacterivory in aquatic environments [21–23]. Models suggest that when phago-mixotrophs are taken into account in trophic network analysis, the transfer of biomass to higher trophic levels increases and mean organism size and sinking carbon fluxes are higher [19]. A recent study [16] reviews the current evidence and importance of phago-mixotrophy in the Arctic ocean where this trophic mode has been documented in chrysophytes (*Ochromonas* spp. and *Dinobryon balticum*), cryptophytes (*Geminigera cryophilia* and *Teleaulax amphioxeia*), prymnesiophytes (*Chrysochromulina spp.*), chlorophytes (*Pyramimonas spp.* and *Micromonas polaris*) as well as a number of dinoflagellates and ciliates species.

The on-going expansion of stratification and nutrient limitation in the Arctic have been associated with an observed increase of the smaller phytoplankton (picophytoplankton: 2-3 *μ*m cell diameter) [24, 25] composed only of eukaryotes since cyanobacteria are nearly absent in polar marine ecosystems [26]. Among the picophytoplankton community, the green alga *M. polaris* [27, 28] dominates in the Arctic ocean [27, 29–32] and its abundance is expected to increase as the stratified oligotrophic areas expand [24, 33, 34]. The physiological plasticity allowing *M. polaris* to dominate the Arctic picoeukaryote community as well as possibly thrive under the climate driven changes observed in the Arctic, is not yet well understood. *M. polaris* was shown in the laboratory, to positively respond to the combination of warming and acidification by increasing its growth rate and biomass production [33]. Phago-mixotrophy would be another advantageous trait that could contribute to the success of *M. polaris* in the Arctic. Under prolonged periods of darkness or low irradiance, phago-mixotrophs could survive, despite reduced or even null rates of photosynthesis, by supplementing their carbon requirements through phagocytosis [16, 35, 36]. Under oligotrophic conditions, phago-mixotrophy can also supply the cell with limiting nutrients [37].

*Micromonas* has been previously hypothesized to be a phago-mixotroph in laboratory and field experiments [38–41]. More than 25 years ago, Gonzales *et al.* [41] reported phago-mixotrophy in a temperate *Micromonas* strain (identified at that time as *M. pusilla*) based on a positive acid lysozyme assay and ingestion of fluorescently labelled bacteria (FLBs) measured by microscopy. More recently, the ability of Arctic pico and nanoplankton microbial communities to consume bacterioplankton has been analyzed by *in situ* experiments using FLBs and yellow-green fluorescent microspheres (YG-beads) as prey. A *Micromonas*-like picoeukaryote, based on its shape and analysis of denaturing gradient gel electrophoresis (DGGE) band sequences, was reported to ingest a significant quantity of prey offered to it [40]. Ingestion of beads was further tested in *M. polaris* strain CCMP2099 under laboratory conditions that compared different light levels and nutrient concentrations. The highest grazing rates were observed under light and low nutrient conditions [38] for which transcriptional response was also investigated [39]. Despite the evidence presented, it is still unclear whether *M. polaris* is capable of ingesting bacteria because of the difficulty to distinguish whether the preys are inside the cells or just externally attached to them [42] using epifluorescence microscopy. Recently, association of YG-beads with *M. polaris* (strain CCMP2099) cells was found after performing feeding experiments with heat-killed cultures [42], suggesting that beads may stick to the surface of the cells resulting in a potential over-estimation of phagocytosis.

In the present paper, we used flow cytometry to analyse prey uptake in several *M. polaris* strains isolated from different regions and depths in the Arctic Ocean, including CCMP2099. We also made predictions of the capacity of *Micromonas* to be a phago-mixotroph from an *in silico* gene-based model [43].

## Materials and Methods

### Strains and culturing conditions

Four *M. polaris* strains and one phago-mixotrophic *Ochromonas triangulata* strain were used in this study. Three of the *M. polaris* strains (RCC2306, RCC4298 and RCC2258) and *O. triangulata* strain RCC21 were obtained from the Roscoff Culture Collection (RCC, http://www.roscoff-culture-collection.org/). The fourth *M. polaris* strain (CCMP2099) was obtained from the Provasoli-Guillard National Center for Culture of Marine Phytoplankton and Microbiota (https://ncma.bigelow.org) and is also available from the RCC as RCC807. The *M. polaris* strains originate from different locations and depths in the Arctic (Table 1). All strains were non-axenic and grown under a 12h:12h light:dark cycle at 80 *μ*E m^−2^ s^−1^ PAR using L1 medium [44] made with artificial sea water (ASW) [45]. All *M. polaris* strains were grown at 4 °C and *O. triangulata* at 20 °C. Cells were acclimated and maintained in mid-exponential growth phase before the beginning of each experiment.

**Table 1.**
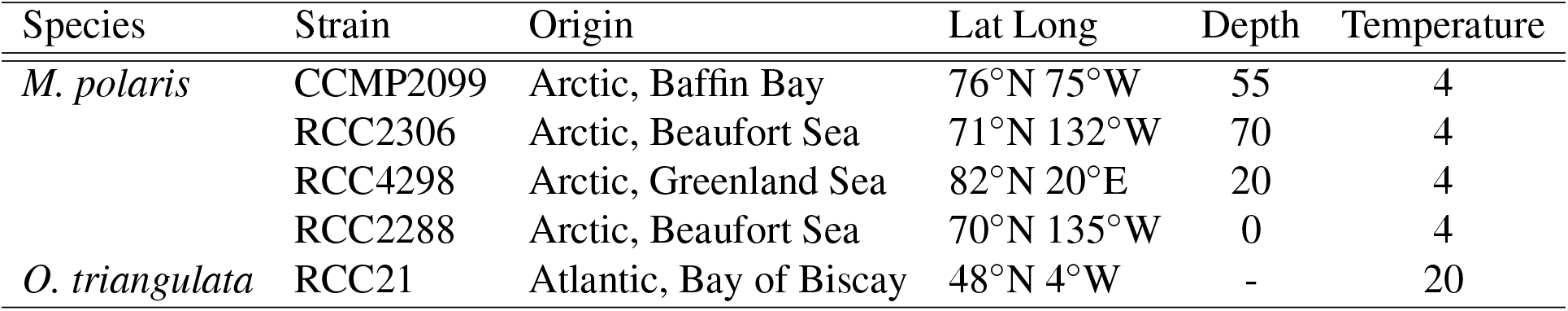
List of algal strains used in this study with isolation region, coordinates, depth (m) and growth temperature (°C).

| Species | Strain | Origin | Lat Long | Depth | Temperature |
| --- | --- | --- | --- | --- | --- |
| *M. polaris* | CCMP2099 | Arctic, Baffin Bay | 76°N 75°W | 55 | 4 |
|  | RCC2306 | Arctic, Beaufort Sea | 71°N 132°W | 70 | 4 |
|  | RCC4298 | Arctic, Greenland Sea | 82°N 20°E | 20 | 4 |
|  | RCC2288 | Arctic, Beaufort Sea | 70°N 135°W | 0 | 4 |
| *O. triangulata* | RCC21 | Atlantic, Bay of Biscay | 48°N 4°W | - | 20 |

### Cell monitoring, feeding estimates and sample fixation

Cells and prey were counted using a Guava easyCyte (Luminex Corporation, USA) flow cytometer (FCM) equipped with a 488 nm laser recording cell counts, forward and side angle light scatter (FALS and SSC), both proxies of cell size, green (525 *±* 30 nm band pass filter) and red (695 *±* 50 nm band pass filter) fluorescence. Cultures under the different experimental conditions were monitored live using red autofluorescence from chlorophyll as a threshold. Flow cytometry was also used to determine the percent of cells with prey (YG-beads and FLBs) in samples fixed using a protocol modified from Sherr *et al.* [46] (acid Lugol’s iodine solution and formaldehyde 3.7%, and cleared with sodium thiosulfate 3%) with a threshold either on red fluorescence or green fluorescence. With the threshold on red fluorescence, cells that contained chlorophyll as well as green fluorescence (same signal as the prey added, YG-beads or FLBs) were considered to be cells containing prey (Figure S1). In addition, to confirm the total concentration of prey added to each flask, the sample was also run with the threshold on green fluorescence. FCM listmodes were analyzed with the Guava easyCyte Suite Software 3.1 (Luminex Corporation, USA).

For each feeding experiment, the ingestion of prey was quantified in each experimental flask by first adding prey and then sub-sampling and fixing after an incubation of 0 (T_0_), 20 (T_20_) and 40 (T_40_) minutes. The T_0_ sample accounts for the physical attachment of prey to the cell and therefore the percent of cells ingesting prey corresponds to the percent of cells with prey at T_20_ or T_40_, minus the percent of cells with prey at T_0_.

### Microscopy

Light-limited *M. polaris* (strain RCC2306) cells were fixed just after the addition of YG-beads (T_0_) with glutaraldehyde (1% final concentration). Fixed cells were sedimented onto formvar coated copper grids for 30 minutes. Grids were then stained with three drops of uranyl acetate 2%, dried and examined using a JEOL JEM1400 transmission electron microscopy (TEM, Jeol, Tokyo, Japan) operating at 80 kV. Images were obtained with a Gatan Orius camera (Roper Scientific SAS, France).

### Major feeding experiments

To test feeding, three different experimental designs were performed with *M. polaris* strains and another fourth with *O. triangulata* (Table S1). Feeding was primarily tested using yellow-green fluorescent polystyrene-based microspheres (YG-beads; 0.5 *μ*m in diameter, Fluoresbrite, Polysciences, Inc., Warrington, PA, USA) as preys. In some cases fluorescently labelled bacteria (FLBs) were used. FLBs were prepared according to the protocol of Sherr *et al.* [47] using the bacteria *Brevundimonas diminuta* (strain CECT313, also named *Pseudomonas diminuta*), obtained from the Spanish Type Culture Collection (CECT, Valencia, Spain).

In experiment type 1 (*M. polaris*-EXP1) feeding was tested for each *M. polaris* strain grown under four different culture conditions. Each treatment was carried out (in duplicates for RCC2306 and RCC4298 and triplicates for RCC2258 and CCMP2099) by transferring a small volume of culture (a few ml in general), previously maintained in mid-exponential growth, to about 40 ml of L1-ASW medium (replete) or ASW without any addition (limited) in a 50 ml culture flask and then placed in the dark or left in the same light conditions as for culture maintenance. Each treatment (light-replete, light-limited, dark-replete and dark-limited) was followed up for 15-17 days and feeding was tested with YG-beads after 7 (Feeding 1) and 14-17 (Feeding 2) days.

Experiment type 2 (*M. polaris*-EXP2) was performed with *M. polaris* strain RCC2306 and RCC2258 and was set-up the same way as EXP1 (in triplicates), but with an additional treatment (light-replete-Ab) in which 1 *μ*l of Penicillin-Streptomycin-Neomycin (PSN) antibiotics solution (Sigma Aldrich P4083) per 1 ml of culture was added at the beginning of the experiment in order to minimize bacteria concentration. Moreover, the five treatments were incubated for only one week and feeding was tested with YG-beads at the end of the incubation (Day 7).

To compare feeding on YG-beads and FLBs, a third type of experiment (*M. polaris*-EXP3) was performed with *M. polaris* RCC4298. For each prey type (YG-beads and FLBs) feeding was tested in triplicate in mid-exponential phase cultures (light-replete).

For all experiments (*M. polaris*-EXP1 to EXP3), the initial concentration for each treatment was 5 x 10^5^ cells ml^−1^. The prey concentration was adjusted in order to achieve a prey to cell ratio of 1.5 to 2.5.

The experimental design of experiments EXP1 and 2 performed with *O. triangulata* was the same and only differed in their replication and number of feeding time points. *O. triangulata*-EXP1 was conducted in duplicate and with three feeding time points (T_0_, T_20_ and T_40_ minutes), and *O. triangulata*-EXP2 in triplicates and two feeding time points (T_0_ and T_40_ minutes). Feeding was tested under two different culture conditions by transferring a small volume of culture, previously maintained in mid-exponential growth, to L1-ASW medium (light-replete) or ASW without any addition (light-limited) and incubated in the same light conditions as for culture maintenance. After one week of incubation, feeding was tested with YG-beads. The third experiment type (*O. triangulata*-EXP3) was performed in parallel with *M. polaris*-EXP3 to compare feeding on YG-beads and FLBs. For each prey type (YG-beads and FLBs) feeding was tested in biological triplicates in mid-exponential phase cultures (light-replete). In a fourth experiment type (*O. triangulata*-EXP4) feeding was tested using FLBs as prey in light-replete culture conditions. EXP4 was performed two times (EXP4a and b) and each time in duplicates.

### Additional experiments

The degree of attachment of YG-beads to cells, immediately after the addition of prey (T_0_) was further examined in a number of additional experiments (*M. polaris*-EXP5) performed with *M. polaris* strains RCC2306 and RCC4298. For *M. polaris* strain RCC2306 the quantification was done in cultures grown under light-replete, light-limited, dark-replete and dark-limited conditions, and for *M. polaris* strain RCC4298 with cultures grown under these four conditions plus light-replete-Ab.

The effects of fixation on the attachment of YG-beads to cells (*M. polaris*-EXP6) was measured by simultaneously comparing feeding in experiments performed with *M. polaris* (strain RCC2306) and run in the flow cytometer live or after fixation with Lugol’s iodine solution and glutaraldehyde (0.25% final concentration). For this experiment, *M. polaris* (strain RCC2306) in mid-exponential (Light-replete) feeding was measured at two time points (T_0_ and T_40_).

Feeding on three different YG-bead sizes (0.5, 1, and 2 *μ*m in diameter) (*M. polaris*-EXP7) was measured in *M. polaris* (strain RCC2306) incubated for one week in light-limited conditions (duplicates). Feeding was measured independently for each bead size using two feeding time points (T_0_ and T_40_).

Changes in the number of cells with YG-beads was measured by continuously running a live sample for 20 minutes immediately after the addition of YG-beads (*M. polaris*-EXP8). Samples were quantified on the FACSCanto (BD Biosciences, USA) flow cytometer with the same configuration as the Guava. For this experiment, cultures of *M. polaris* (strain RCC2306), previously incubated for one week in light-limited conditions, were used. Two ratios of Beads to cells were tested (ratio of 1:1 and 2:1) in duplicates.

The percent of cells potentially containing food vacuoles (EXP9) was quantified in *M. polaris* (strain RCC2306) and *O. triangulata*, stained with the probe LysoSensor Green DND-189 (Thermo Fisher Scientific), that accumulates in acid cellular compartments like food vacuoles, at a final concentration of 1 *μ*M. After the addition of Lysosensor, cells were incubated in the dark for 8 minutes and measured for 2 minutes using the Guava easyCyte (Luminex Corporation, USA) flow cytometer (triggered on green fluorescence). Cells with higher green fluorescence, after incubation with Lysosensor, were considered as potentially containing food vacuoles (Figure S2). The cells used for this test came from light-limited cultures (duplicates), from *O. triangulata*-EXP1 and *M. polaris*-EXP2, on which feeding experiments were performed.

### Data analysis

Data processing, graphics and statistical analyses were performed using the R software [48] using in particular the package set *tidyverse*. Pairwise comparisons were performed with the *t.test* function to calculate p-values based on Student (assuming equal variances) and Welch (assuming unequal variances) t-test.

### Trophic mode predictions from genome and transcriptome analysis

Predicted peptides from whole genome [49, 50] or transcriptome data [51] were downloaded from publicly available databases as detailed in Table S2. Because computational predictions are based on presence/absence information, predicted peptides from independent transcriptome assemblies of the same strain were concatenated to include as much information about each strain as possible. Computational prediction of phagocytotic, photosynthetic, and prototrophic capabilities were completed as in Burns *et al.* [43]. This involved scoring a set of 14,095 protein profile hidden Markov models (HMMs) that were derived by clustering all proteins in 35 reference eukaryote genomes of known trophic mode against all proteins from each test genome or transcriptome. HMM profiles with a full sequence e-value ≤ 10^−5^ and a single domain e-value *≤* 10^−4^ to any protein in a test genome or transcriptome were marked as “present” for that organism. Predictive models of trophic modes were built by grouping the reference eukaryotes by known trophic mode and discovering HMMs from the set of 14,095 that had differential presence/absence patterns between groups. Those HMMs whose presence/absence patterns differed according to trophic mode were annotated against SwissProt and grouped by gene ontology (GO) terms. GO categories were scored per reference organism and a best predictive set of GO terms was selected for each trophic mode using machine learning algorithms, forming the core of the predictive trophic mode models. Each test genome/transcriptome was scored for the predictive GO categories of the trophic mode models using its HMM presence/absence vector. Final prediction probabilities for each test genome/transcriptome were calculated against the reference trophic mode models using a probability neural network. To visualize the prediction output, which exists in four dimensions with three degrees of freedom (phagocytosis, photosynthesis, prototrophy, and a fourth dependent dimension for absence of each trophic mode), predictions were normalized such that the sum of the three predictions plus the probability of not fitting each trophic mode equaled 1 using the relation: 1 − (*p_phago_* + *p_proto_* + *p_photo_*)/3. The fractional independent probabilities of each trophic mode and the dependent absence number were mapped to 4-dimensional color space and projected onto a circle using scripts adapted from the R package “pavo” [52]. Scripts are available at https://github.com/burnsajohn/predictTrophicMode.

## Results

### Feeding experiments

*M. polaris* feeding was analyzed in four strains (CCMP2099, RCC2306, RCC4298 and RCC2258) (Table 1) using a slight modification of the protocol described in Sherr *et al.* (1993) [46]. We determined the percentage of cells feeding on YG-beads or FLBs using flow cytometry to quantify the proportion of cells with prey. Compared to epifluorescence microscopy, which is low throughput allowing examination of at most 100 to 200 cells per sample, flow cytometry allows screening of a large number of cells per sample (typically several thousand). It also circumvents ambiguities that arise with microscopy when cells and prey randomly overlap during the filtration process [42]. To validate our approach we used the phago-mixotrophic *O. triangulata* strain RCC21 as a positive control.

*M. polaris* strains were grown under a combination of 2 light regimes and 2 nutrient concentrations (*M. polaris*-EXP1: light-nutrient replete, light-nutrient limited, dark-nutrient replete and dark-nutrient limited) and experiments took place over a period of 15 to 17 days. Feeding was examined with YG-beads after 7 (Feeding 1) and 14-17 days (Feeding 2). Clear negative growth effects under darkness and nutrient limitation conditions were observed for all strains. Overall, for all 4 strains under dark conditions, growth ceased between day 4 and day 7 and thereafter cell concentration remained stable (Figure 1). For cultures grown under low nutrient conditions (nutrient-limited), a decrease in growth rate was observed after one week of incubation (Figure 1). Additional signs of the effect of darkness and nutrient limitation were observed in FALS (proxy of cell size) and chlorophyll fluorescence: for example FALS decreased for cells in the dark (Figure S3). In all feeding experiments we observed that cells at time T_0_, immediately following addition of the beads, already had a number of beads associated with them. However, no significant difference was observed between the percent of cells with YG-beads at T_0_ and T_20_ or T_40_ whatever the growth phase or the culture condition (Figure 2 and Table S3).

**Figure 1.**
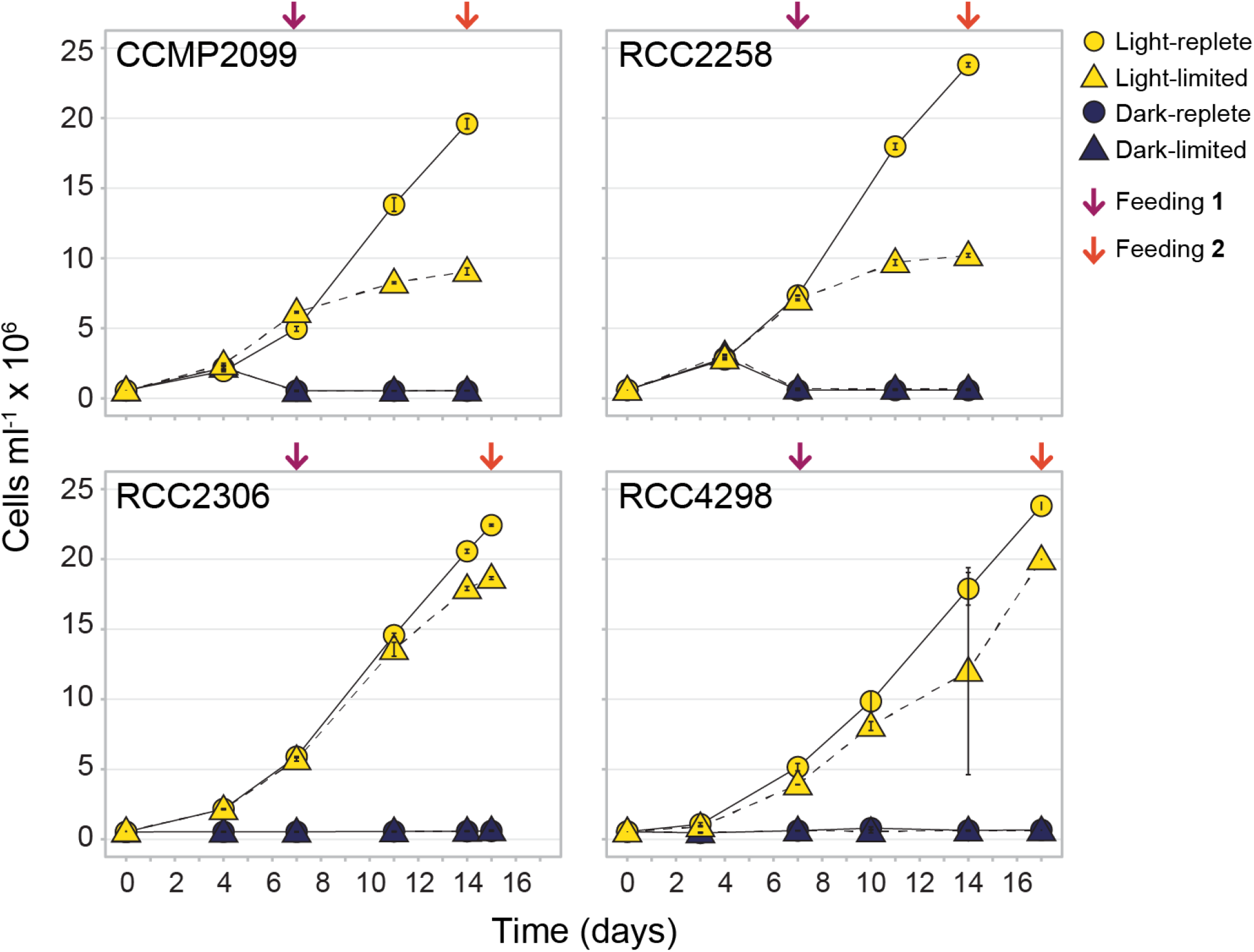
Growth curves for each *M. polaris* strain grown under four treatments (*M. polaris*-EXP1). Arrows indicate the time point (days) when a feeding experiment was performed. Error bars correspond to standard deviation and in some cases are smaller than the symbol used.

**Figure 2.**
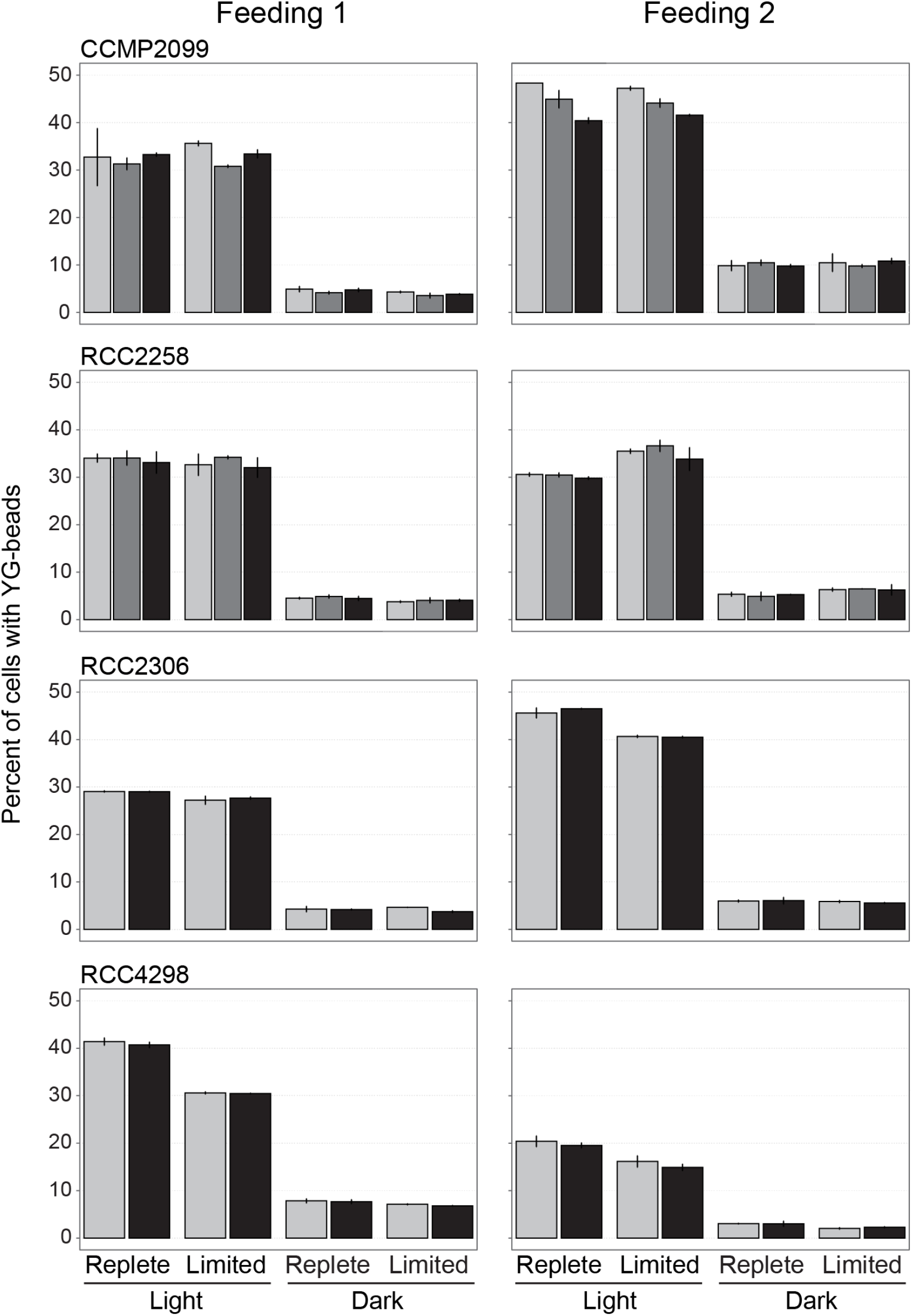
Percent of *M. polaris* cells with YG-beads (*M. polaris*-EXP1) for each strain and treatment at each feeding experiment. The color of the bars represent the time point (in minutes) after the addition of YG-beads (0 minutes; light grey, 20 minutes; dark grey, 40 minutes; black). Error bars correspond to standard deviation.

We questioned whether the absence of feeding on YG-beads could have been due to the presence of bacteria in the cultures. In order to address this issue, we performed a second series of experiments in which we included a fifth condition by adding antibiotics to a light-nutrient replete culture (*M. polaris*-EXP2). This was only performed with two of the *M. polaris* strains (RCC2258 and RCC2306) and a single feeding experiment was conducted after one week. No feeding was detected under any of the culture conditions (Figure S4).

We then compared feeding on YG-beads vs. FLBs as prey (*M. polaris*-EXP3) since the prey type may influence feeding behavior. Again no feeding was observed when using either YG-beads or FLBs (Table 2). In contrast, in the four experiments performed with *O. triangulata* (EXP1 to EXP4) we observed feeding on YG-beads and FLBs that ranged from 7 to 14 and 21 to 27 percent of cells feeding on each prey type respectively, suggesting that *Ochromonas* clearly preferred FLBs over YG-beads (Table 2 and S3).

**Table 2.**
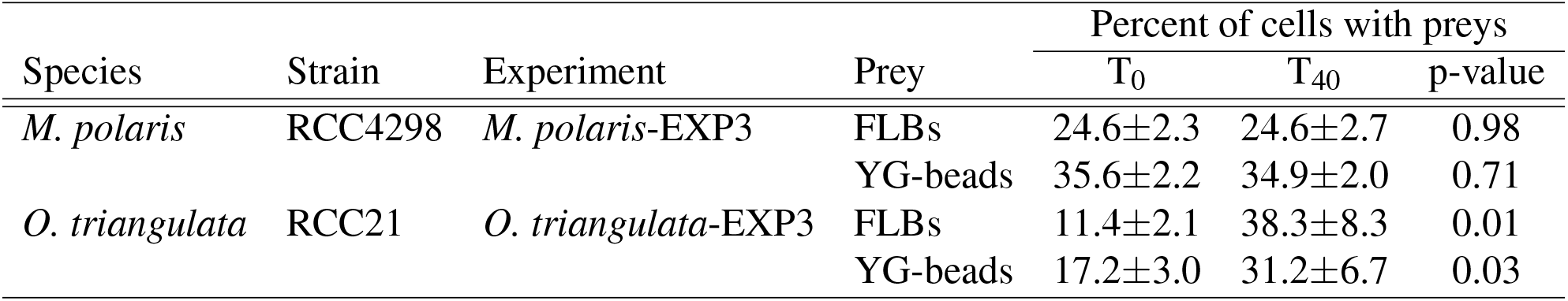
*M. polaris*-EXP3. Comparison between preys (YG-beads and FLBs) in feeding experiments. The percent of cells with preys (Mean±sd) is indicated for each time point after the addition of prey (T_0_ and T_40_, where the number corresponds to minutes). The p-value of the difference between T_0_ and T_40_ time points is also indicated.

The percentage of cells with 0.5 *μ*m YG-beads at T_0_ was clearly related to cell concentration (Figure 3) and saturated at high cell concentrations to roughly 50%. This relationship was best represented by a log-linear relationship (R^2^ = 0.79). Cell size did not seem to have an influence since *Micromonas* (≃ 1.5 *μ*m) and *Ochromonas* (≃ 5 *μ*m) fit the same curve. The number of cells with YG-beads did not change over time as demonstrated by monitoring live cells of *M. polaris* (strain RCC2306) in the presence of YG-beads by flow cytometry over a 20 minutes period (Figure S5).

**Figure 3.**
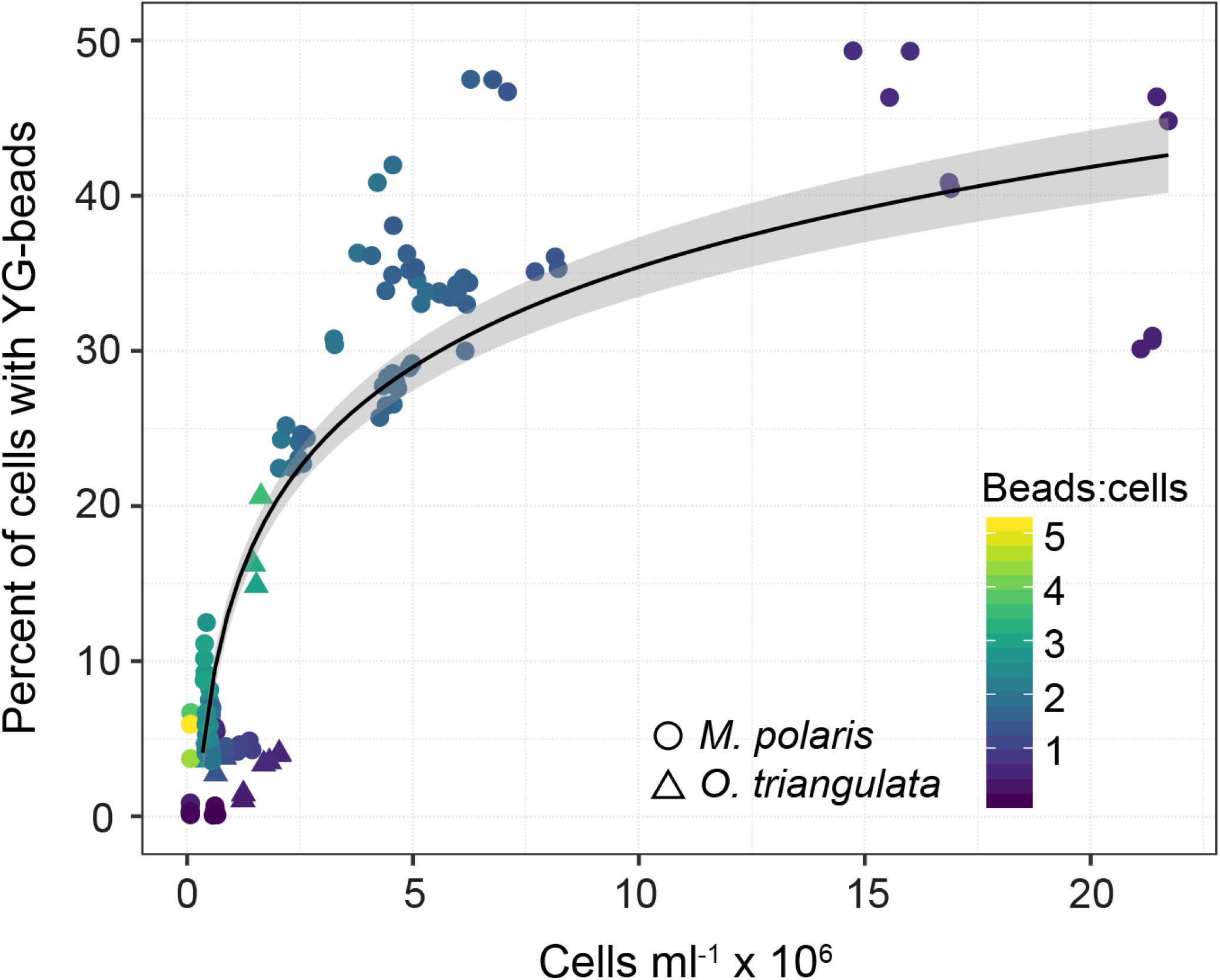
Relationship between percent of cells with YG-beads and cell concentration immediately after the addition of YG-beads (T_0_). The line and spread represents a fitted log-linear relationship (R^2^ = 0.79). Circles correspond to *M. polaris* and triangles to *O. triangulata* experiments.

The association of beads and cells did not seem to be linked to bead size. We still observed association of 1 and 2 *μ*m YG-beads with cells at T_0_, even though the 2 *μ*m beads are close in size to *M. polaris* cells and no differences were observed between the percent of cells with YG-beads at T_0_ and T_40_ (Figure S6 and Table S4). Fixation does not seem to impact the association of beads at T_0_ as we observed this co-association when samples were run live or fixed with Lugol’s solution or glutaraldehyde (Table S5). External attachment of YG-beads to cells of *M. polaris* (strain RCC2306) was visualized by TEM (Figure S7).

Since phagotrophy in *M. polaris* had been proposed previously based on an observation of acidic food vacuoles [42, 53], *M. polaris* light-nutrient limited cultures (from EXP-2, which did not feed on YG-beads) were stained with the acidotropic LysoSensor fluorescent probe. No significant difference was observed in green LysoSensor fluorescence between unstained and stained cells, whereas for *O. triangulata* (EXP1 light-nutrient limited) green fluorescence increased 3.5 times after staining, suggesting the presence of food vacuoles in the latter species (Figure S2 and Table S6).

### Trophic mode predictions

Phagocytotic, photosynthetic, and prototrophic capacity of protists can be predicted based on their genome or transcriptome composition [43]. We used this approach to analyze gene composition of a number of microalgae including *Micromonas* (Table S2). The predictions confirm that known phago-mixotrophs like *Dinobryon sp.*, Pedinelalles sp., *Ochromonas triangulata*, and *Prymnesium parvum* have and express a battery of genes consistent with their observed lifestyle coherent with the capacity for phagocytosis, photosynthesis, and prototrophy (Figure 4). Presumed photo-autotrophs like members of the genus *Ostreococcus* lack genes consistent with the capacity for phagocytosis, but have genes consistent with the capacity for photosynthesis and prototrophy (Figure 4). Similarly, all members of the genus *Micromonas* are predicted to be photo-autotrophs as they contain genes consistent with photosynthesis and prototrophy, but lack genes consistent with the capacity for phagocytosis (Figure 4).

**Figure 4.**
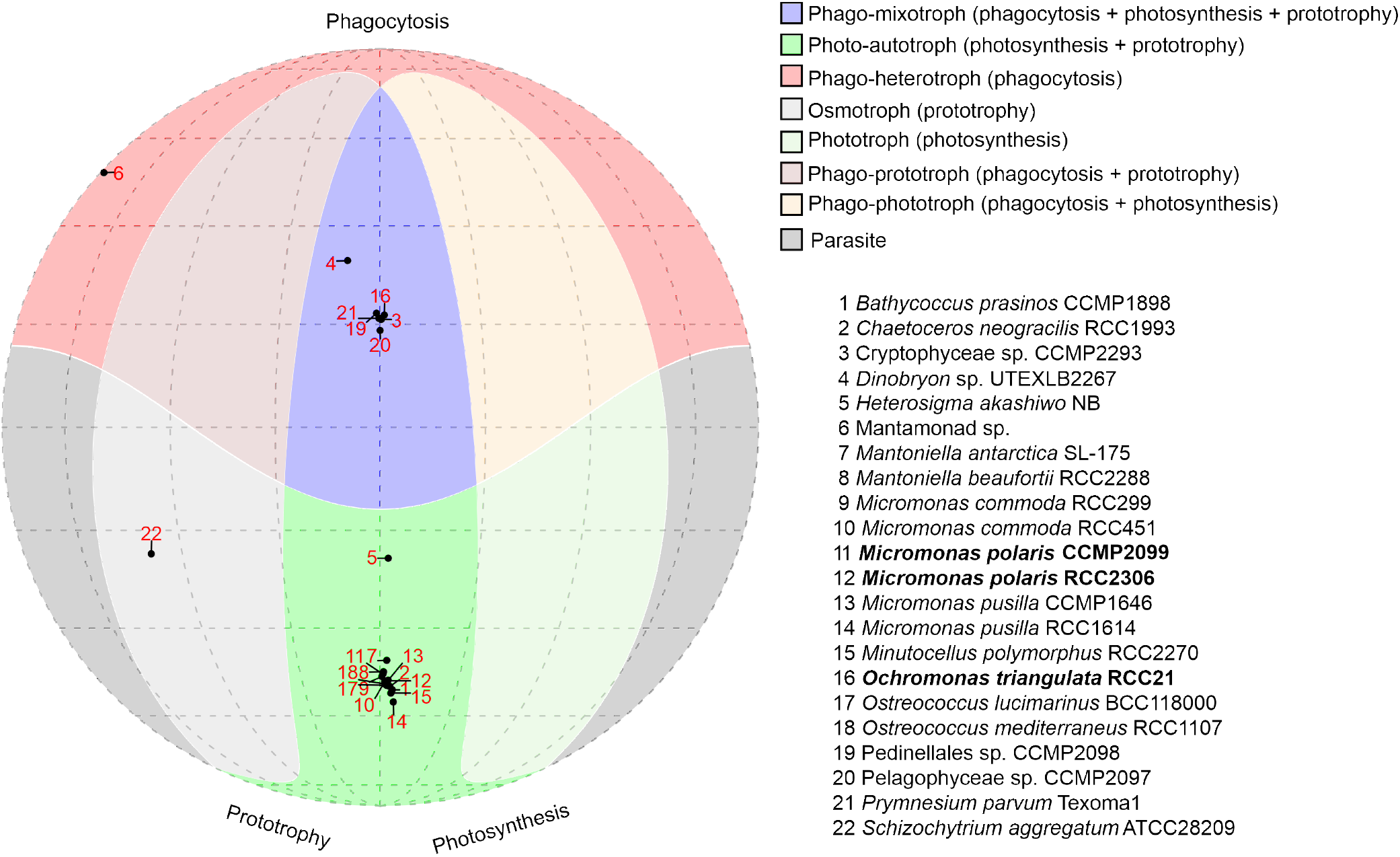
Trophic mode predictions from genome and transcriptome analysis. Predictions in three dimensions, phagocytosis, photosynthesis, and prototrophy were projected onto the circle. Shaded regions indicate regions where 0 (Parasite, gray, lower edges) to all 3 (Phag-mixotroph, blue, upper central region) predictions cross the positive threshold of 50% probability of a strain possesing a given function. Organisms positive for phagotrophy will be in the upper hemishpere. Organisms positive for photosynthesis will be in the middle to right region. Organisms positive for prototrophy will be in the middle to left region. Organisms negative for all predictions will be in the lower gray edge regions. Strains in bold correspond to those used in the feeding experiments.

## Discussion

We examined feeding of *M. polaris* on preys in a series of experiments with four different strains (CCMP2099, RCC2306, RCC4298 and RCC2258) grown under different light and nutrient conditions using flow cytometry to monitor prey uptake (Table S1). In none of these experiments (Figure 2 and Table S3), significant differences were detected between the number of *M. polaris* cells associated with preys at T_0_ and other time points (T_20_ or T_40_). We also tested different fixation methods vs. live cells and three different diameters of beads (0.5, 1, and 2 *μ*m in diameter) without detecting any clue of active uptake by *M. polaris*. No evidence of phago-mixotrophy was found when using the acidotropic LysoSensor dye in *M. polaris* light-nutrient limited cultures. Trait-based computational analysis of available genomes and transcriptomes confirmed that *Micromonas* lack genes consistent with the capacity for phagocytosis. These data are in contrast to what was observed for the known phagotroph *O. triangulata* (strain RCC21) that always displayed evidence of prey uptake when using similar approaches to the one we used for *M. polaris* strains and fits the computational profile of a phago-mixotroph. None of our evidence is consistent with the consideration of *M. polaris* as a phago-mixotroph.

In each of our experiments, there was a considerable number of *M. polaris* cells at T_0_ associated with prey, immediately following addition of prey to the cultures, before time had elapsed for prey ingestion. The percentage of cells with 0.5 *μ*m YG-beads at T_0_ appears to be log-linearly related to the culture concentration (Figure 3), suggesting the association is the result of a physical property of the cells surface rather than an active behavior that the cells execute. The external attachment of YG-beads to *M. polaris* cells was also visualized by electron microscopy (Figure S7). Such passive associations of cells with beads have recently been observed by Wilken *et al.* (2019) [42] using flow cytometry. They observed that the proportion of cells associated with beads at T_0_ was much larger for heat-killed vs. live cells and that it increased with time for cultures left in the dark. Such external attachment of particles or bacterial cells to phytoplankton cell surfaces may be enhanced by phycosphere properties [54, 55] which mainly consist of polysaccharides released by the cells [56–58]. The “stickiness” properties of abundant exopolysaccharides have mainly been studied in diatoms [55, 59, 60]. Bacteria colonization of this sticky phycosphere is a function of the probability of random encounters of phytoplankton and bacteria which is influenced by both cell concentration and cell motility [55].

Our experimental conditions were very similar to those reported by McKie-Krisberg *et al.* (2014) [38]. We used the same *M. polaris* strain (CCMP2099), dark and light conditions, ASW as medium, 0.5 *μ*m beads, Lugol’s iodine fixation and short term incubation (40 min.). The main methodological difference is that we used flow cytometry analysis of cell suspensions instead of epifluorescence microscopy of filtered samples. Our approach has many advantages over epifluorescence microscopy: it allows counting of a much larger number of cells (typically several thousand vs. 100), it is faster and not operator dependent resulting in less potential biases related to individual operator interpretation and with food particles randomly overlapping with cells during filtration. The latter problem is demonstrated in McKie-Krisberg *et al.* (2014) [38]: their differential interference contrast and confocal microscopy images (Figure 1 c-d in [38]) aimed at demonstrating a YG-bead inside a *M. polaris* cell are inconclusive as the bead is at the edge of the cell (probably externally attached) which closely resembles the TEM images obtained in the present study. The two other papers that have reported phago-mixotrophy in *Micromonas* [40, 41] may have suffered from the same problem, i.e. attachment of preys to cells. Moreover in the Sanders *et al.* (2012) [40] paper on natural communities the identity of the potential grazer was only “tentatively identified as *Micromonas*” from the presence of a DGGE band with a *Micromonas* sequence. A study that examined gene expression of *M. polaris* strain CCMP2099 under nutrient stress conditions that reportedly influence grazing rate failed to find differential expression of any gene linked to the process of phagocytosis in *M. polaris* [39]. The authors propose that *M. polaris* may constitutively express phagocytosis proteins to support low-level grazing. However, a study on the model phagocyte *Dictyostelium discoideum* suggests that an increase in phagocytosis can indeed be linked to differential expression of genes involved in the process [61]. An alternative hypothesis regarding the gene expression results from *M. polaris*, supported by the data presented here, is that members of the genus *Micromonas* are not phagocytotic and therefore have no mechanism for differential expression of genes linked to phagocytosis. Sets of proteins identified as part of the phagosome compartment are broadly distributed among phagocyte and non-phagocyte organisms and only a small subset of those proteins are indicative that a species has the capacity for phagocytosis [62]. Computational models show that members of the genus *Micromonas* lack those indicative proteins, reinforcing our hypothesis that *Micromonas* is not a phago-mixotroph.

Bacterial phagocytosis has been found everywhere across the eukaryotic tree of life [20], but most laboratory studies on phago-mixotophy have focused on a few species such as the chrysophyte *Ochromonas* sp. (e.g. [63–65]), the haptophytes *Prymnesium parvum* (e.g. [66, 67]) and *Chrysochromulina* spp. [68, 69], and dinoflagellates (e.g. [70–72]). Among green algae in addition to the works on *Micromonas* mentioned previously, only a few studies have been performed with 6 other species described as phago-mixotrophs (*Pyramimonas gelicola* [73], *Pyramimonas tychotreta* and *Mantoniella antarctica* [74], *Cymbomonas tetramitiformis* [75, 76], *Nephroselmis rotunda* and *Nephroselmis pyriformis* [77]). None of these species fall however in the picoplankton size range. Interestingly none of the green algae (in addition to *Micromonas*) for which the trait-based computational analysis was performed (*Bathycoccus*, *Ostreococcus*, *Mantoniella*, including *M. antarctica*) showed evidence for phago-mixotrophy.

It is now acknowledged that phago-mixotrophy is a widespread trait in planktonic communities and has profound implications for marine ecosystem functioning [37, 78]. In particular phago-mixotrophy is believed to provide a competitive advantage to photosynthetic organisms under otherwise limiting environmental conditions (e.g. low light, low nutrients). Based on our results and in contrast to what has been suggested previously, this seems less true for pico-size eukaryotes of the green lineage. Despite being primary players in resource limited environments, green picoeukaryotes, notably abundant members of the Mamiellophyceae such as *Micromonas* and *Bathycoccus*, are likely to rely on other strategies to thrive in oligotrophic waters [79] or survive through polar winter [80, 81].

The evidence presented in this paper indicating that *M. polaris* is unlikely to be phago-mixotroph has profound impacts in present and future predictions of Arctic primary production, because of the importance and predicted increasing concentrations of this species in the Arctic Ocean [24]. If indeed *M. polaris* is not a phago-mixotroph, the question of how it survives during the long Arctic winter and how it is able to develop during the Spring bloom that starts with very low light condition under the snow-covered ice [82] remains open.

## Acknowledgments

This work was supported by ANR contract PhytoPol (ANR-15-CE02-0007). We thank Robert Sanders and Rebecca Gast for hosting VJ and training her on techniques to determine mixotrophy in phytoplankton. We also thank Dominique Marie for help with flow cytometry, Christian Jeanthon for help and advice with fluorescently labelled bacteria, Sophie Le Panse from the Merimage microscopy platform at the Roscoff Marine Station for assistance with the transmission electron micrographs and the Roscoff Culture Collection for providing of the algal strains.

## Author contributions statement

DV and VJ conceived the study. VJ and FLG collected and processed the samples. VJ, JB and DV analyzed the data. VJ, JB and DV drafted the manuscript. VJ, DV, FLG, FN and JB edited the final version of the paper.

## Additional information

## Data availability

Protocols are available at protocols.io at https://www.protocols.io/edit/mixotrophy-quantification-of-the-percent-of-phytop-be2vjge6/steps. Scripts for trophic mode prediction and visualization are available at https://github.com/burnsajohn/predictTrophicMode.

## Competing interests

The authors declare no competing interests.

## Supplementary Material

**Table S1.** Experimental scheme. Replete correspond to cultures grown in Artifical Sea Water (ASW) with L1 medium components added and limited to cultures grown in ASW without any addition. Ab correspond to cultures for which 1 *μ*l of Penicillin-Streptomycin-Neomycin (PSN) antibiotics solution was added to 1 ml of culture. The time points on which the percent of cells with prey was measured is indicated (T_0_, T_20_ and T_40_, where the subscript corresponds to minutes). LR: Light nutrient replete, LL: Light nutrient limited, DR: Dark nutrient replete, DL: Dark nutrient limited, LR-Ab: Light nutrient replete with antibiotics.

| Code | Conditions | Preys | Genus | Time points (mins) |
| --- | --- | --- | --- | --- |
| EXP1 | LR, LL, DR, DL | YG-beads | *Micromonas*, *Ochromonas* | 0, 20, 40 |
| EXP2 | LR, LL, DR, DL, LR-Ab | YG-beads | *Micromonas*, *Ochromonas* | 0, 20, 40 |
| EXP3 | LR | YG-beads, FLBs | *Micromonas*, *Ochromonas* | 0, 40 |
| EXP4 | LR | FLBs | *Ochromonas* | 0, 40 |
| EXP5 | LR, LL, DR, DL, LR-Ab | YG-beads | *Micromonas* | 0 |
| EXP6 | LR | YG-beads | *Micromonas* | 0, 40 |
| EXP7 | LL | YG-beads | *Micromonas* | 0, 40 |
| EXP8 | LR | YG-beads | *Micromonas* | 0 to 20 |
| EXP9 | LL | YG-beads | *Micromonas*, *Ochromonas* |  |

**Table S2.** List of strains used for transcriptome analysis. MMETSP corresponds to the Marine Microbial Eukaryote Transcriptome Sequencing Project [51]. METDB corresponds to the micro-eukaryotic marine species transcriptomes database available from http://metdb.sb-roscoff.fr/metdb/.

| Sequence Source | Species | Strain | Database | Reference |
| --- | --- | --- | --- | --- |
| Transcriptome | *Bathycoccus prasinus* | CCMP1898 | MMETSP | MMETSP1399 |
| Transcriptome | *Chaetoceros neogracilis* | RCC1993 | MMETSP | MMETSP1336 |
| Transcriptome | Cryptophyceae sp. | CCMP2293 | MMETSP | MMETSP0986 |
| Transcriptome | Cryptophyceae sp. | CCMP2293 | MMETSP | MMETSP0987 |
| Transcriptome | Cryptophyceae sp. | CCMP2293 | MMETSP | MMETSP0988 |
| Transcriptome | Cryptophyceae sp. | CCMP2293 | MMETSP | MMETSP0989 |
| Transcriptome | *Dinobryon* sp. | UTEXLB2267 | MMETSP | MMETSP0019 |
| Transcriptome | *Dinobryon* sp. | UTEXLB2267 | MMETSP | MMETSP0020 |
| Transcriptome | *Dinobryon* sp. | UTEXLB2267 | MMETSP | MMETSP0812 |
| Transcriptome | *Heterosigma akashiwo* | NB | MMETSP | MMETSP0416 |
| Transcriptome | *Mantoniella antarctica* | SL-175 | MMETSP | MMETSP1106 |
| Transcriptome | *Mantoniella beaufortii* | RCC2288 | MMETSP | MMETSP1326 |
| Genome | *Micromonas commoda* | RCC299 | NCBI | GCF<sub>0</sub>00090985.2 |
| Transcriptome | *Micromonas commoda* | RCC451 | MMETSP | MMETSP1400 |
| Transcriptome | *Micromonas polaris* | CCMP2099 | MMETSP | MMETSP0802 |
| Transcriptome | *Micromonas polaris* | CCMP2099 | MMETSP | MMETSP1390 |
| Transcriptome | *Micromonas polaris* | RCC2306 | MMETSP | MMETSP1327 |
| Transcriptome | *Micromonas pusilla* | CCMP1646 | MMETSP | MMETSP1080 |
| Transcriptome | *Micromonas pusilla* | RCC1614 | MMETSP | MMETSP1402 |
| Transcriptome | *Minutocellus polymorphus* | RCC2270 | MMETSP | MMETSP1322 |
| Transcriptome | *Ochromonas triangulata* | RCC21 | METDB | METDB-00278 |
| Transcriptome | *Ostreococcus lucimarinus* | BCC118000 | MMETSP | MMETSP0939 |
| Transcriptome | *Ostreococcus mediterraneus* | RCC1107 | MMETSP | MMETSP0938 |
| Transcriptome | Pedinellales sp. | CCMP2098 | MMETSP | MMETSP0990 |
| Transcriptome | Pedinellales sp. | CCMP2098 | MMETSP | MMETSP0991 |
| Transcriptome | Pedinellales sp. | CCMP2098 | MMETSP | MMETSP0992 |
| Transcriptome | Pedinellales sp. | CCMP2098 | MMETSP | MMETSP0993 |
| Transcriptome | Pelagophyceae sp. | CCMP2097 | MMETSP | MMETSP0974 |
| Transcriptome | Pelagophyceae sp. | CCMP2097 | MMETSP | MMETSP0975 |
| Transcriptome | Pelagophyceae sp. | CCMP2097 | MMETSP | MMETSP0976 |
| Transcriptome | Pelagophyceae sp. | CCMP2097 | MMETSP | MMETSP0977 |
| Transcriptome | *Prymnesium parvum* | Texoma1 | MMETSP | MMETSP0006 |
| Transcriptome | *Prymnesium parvum* | Texoma1 | MMETSP | MMETSP0007 |
| Transcriptome | *Prymnesium parvum* | Texoma1 | MMETSP | MMETSP0008 |
| Transcriptome | *Prymnesium parvum* | Texoma1 | MMETSP | MMETSP0814 |
| Transcriptome | *Prymnesium parvum* | Texoma1 | MMETSP | MMETSP0815 |
| Transcriptome | *Prymnesium parvum* | Texoma1 | MMETSP | MMETSP1083 |
| Transcriptome | *Schizochytrium aggregatum* | ATCC28209 | MMETSP | MMETSP0965 |

**Table S3.** Summary of experimental conditions and results for all experiment performed with *M. Polaris* and *O. triangulata* strains. The percent of cells with preys (Mean±sd) is indicated for each time point after the addition of prey (T_0_, T_20_ and T_40_, where the subscript corresponds to minutes). The last four columns correspond to Student and Welsh p-values.

| Species | strain | EXP type | Treatment | Prey | Feeding | Replication | Ratio prey to cells | T<sub>0</sub> | T<sub>20</sub> | T<sub>40</sub> | T<sub>40</sub>-T<sub>0</sub> | Student T<sub>0</sub> vs. T<sub>20</sub> | Welch T<sub>0</sub> vs. T<sub>20</sub> | Student T<sub>0</sub> vs. T<sub>40</sub> | Welch T<sub>0</sub> vs. T<sub>40</sub> |
| --- | --- | --- | --- | --- | --- | --- | --- | --- | --- | --- | --- | --- | --- | --- | --- |
| *Ochromonas triangulata* | RCC21 | EXP1 | Light-replete | YG-beads | 1 | 2 | 0.6 | 4.0±0.1 | 9.3±0.3 | 12.3±0.3 | 8.3 | 0.001 | 0.016 | 0.001 | 0.011 |
| *Ochromonas triangulata* | RCC21 | EXP1 | Light-limited | YG-beads | 1 | 2 | 0.6 | 3.4±0.1 | 7.6±0.0 | 9.9±0.4 | 6.5 | 0.000 | 0.011 | 0.002 | 0.016 |
| *Ochromonas triangulata* | RCC21 | EXP2 | Light-replete | YG-beads | 1 | 2 | 1.3 | 3.3±0.8 |  | 9.4±2.2 | 6.2 |  |  | 0.066 | 0.130 |
| *Ochromonas triangulata* | RCC21 | EXP2 | Light-limited | YG-beads | 1 | 2 | 2.3 | 3.8±0.2 |  | 15.7±0.5 | 11.9 |  |  | 0.001 | 0.006 |
| *Ochromonas triangulata* | RCC21 | EXP3 | Light-replete | FLBs | 1 | 3 | 1.6 | 11.4±2.1 |  | 38.3±8.3 | 26.9 |  |  | 0.006 | 0.025 |
| *Ochromonas triangulata* | RCC21 | EXP3 | Light-replete | YG-Beads | 1 | 3 | 3.2 | 17.2±3.0 |  | 31.2±6.7 | 14.0 |  |  | 0.029 | 0.050 |
| *Ochromonas triangulata* | RCC21 | EXP4a | Light-replete | FLBs | 1 | 2 | 2.1 | 4.6±0.1 |  | 25.1±0.1 | 20.5 |  |  | 0.000 | 0.000 |
| *Ochromonas triangulata* | RCC21 | EXP4b | Light-replete | FLBs | 1 | 2 | 2.8 | 7.2±0.8 |  | 33.4±1.4 | 26.2 |  |  | 0.002 | 0.006 |
| *Micromonas polaris* | CCMP2099 | EXP1 | Dark-replete | YG-beads | 1 | 3 | 2.3 | 4.9±0.6 | 4.1±0.3 | 4.7±0.4 | -0.2 | 0.134 | 0.154 | 0.707 | 0.710 |
| *Micromonas polaris* | CCMP2099 | EXP1 | Dark-limited | YG-beads | 1 | 3 | 2.2 | 4.3±0.2 | 3.5±0.6 | 3.8±0.2 | -0.5 | 0.092 | 0.126 | 0.044 | 0.047 |
| *Micromonas polaris* | CCMP2099 | EXP1 | Light-replete | YG-beads | 1 | 3 | 1.7 | 32.7±6.1 | 31.3±1.3 | 33.3±0.4 | 0.5 | 0.712 | 0.727 | 0.886 | 0.893 |
| *Micromonas polaris* | CCMP2099 | EXP1 | Light-limited | YG-beads | 1 | 3 | 1.5 | 35.6±0.6 | 30.8±0.3 | 33.4±0.9 | -2.2 | 0.000 | 0.001 | 0.024 | 0.032 |
| *Micromonas polaris* | CCMP2099 | EXP1 | Dark-replete | YG-beads | 2 | 3 | 2.8 | 9.8±1.1 | 10.5±0.6 | 9.8±0.3 | -0.1 | 0.451 | 0.464 | 0.922 | 0.925 |
| *Micromonas polaris* | CCMP2099 | EXP1 | Dark-limited | YG-beads | 2 | 3 | 2.8 | 10.5±1.9 | 9.8±0.3 | 10.8±0.6 | 0.3 | 0.551 | 0.579 | 0.775 | 0.784 |
| *Micromonas polaris* | CCMP2099 | EXP1 | Light-replete | YG-beads | 2 | 3 | 0.7 | 48.3±1.7 | 44.9±1.9 | 40.4±0.6 | -7.9 | 0.082 | 0.083 | 0.002 | 0.009 |
| *Micromonas polaris* | CCMP2099 | EXP1 | Light-limited | YG-beads | 2 | 3 | 1.5 | 47.2±0.5 | 44.1±0.9 | 41.6±0.2 | -5.7 | 0.007 | 0.015 | 0.000 | 0.000 |
| *Micromonas polaris* | RCC2258 | EXP1 | Dark-replete | YG-beads | 1 | 3 | 2.1 | 4.5±0.2 | 4.9±0.4 | 4.5±0.5 | -0.0 | 0.242 | 0.255 | 0.899 | 0.901 |
| *Micromonas polaris* | RCC2258 | EXP1 | Dark-limited | YG-beads | 1 | 3 | 2.0 | 3.8±0.2 | 4.1±0.6 | 4.1±0.3 | 0.3 | 0.458 | 0.479 | 0.219 | 0.221 |
| *Micromonas polaris* | RCC2258 | EXP1 | Light-replete | YG-beads | 1 | 3 | 1.6 | 34.0±0.9 | 34.1±1.6 | 33.1±2.3 | -0.9 | 0.979 | 0.979 | 0.557 | 0.574 |
| *Micromonas polaris* | RCC2258 | EXP1 | Light-limited | YG-beads | 1 | 3 | 1.6 | 32.6±2.3 | 34.2±0.4 | 32.0±2.1 | -0.6 | 0.315 | 0.365 | 0.756 | 0.757 |
| *Micromonas polaris* | RCC2258 | EXP1 | Dark-replete | YG-beads | 2 | 3 | 2.4 | 5.3±0.5 | 4.9±0.9 | 5.3±0.1 | -0.1 | 0.499 | 0.513 | 0.849 | 0.816 |
| *Micromonas polaris* | RCC2258 | EXP1 | Dark-limited | YG-beads | 2 | 3 | 2.6 | 6.3±0.4 | 6.5±0.0 | 6.3±1.1 | -0.1 | 0.522 | 0.555 | 0.936 | 0.938 |
| *Micromonas polaris* | RCC2258 | EXP1 | Light-replete | YG-beads | 2 | 3 | 0.5 | 30.6±0.4 | 30.5±0.5 | 29.8±0.3 | -0.8 | 0.795 | 0.795 | 0.065 | 0.069 |
| *Micromonas polaris* | RCC2258 | EXP1 | Light-limited | YG-beads | 2 | 3 | 1.4 | 35.5±0.5 | 36.6±1.2 | 33.8±2.4 | -1.7 | 0.211 | 0.244 | 0.313 | 0.360 |
| *Micromonas polaris* | RCC2306 | EXP1 | Dark-replete | YG-beads | 1 | 2 | 2.6 | 4.3±0.6 |  | 4.2±0.2 | -0.1 |  |  | 0.853 | 0.865 |
| *Micromonas polaris* | RCC2306 | EXP1 | Dark-limited | YG-beads | 1 | 2 | 2.5 | 4.6±0.0 |  | 3.7±0.2 | -0.9 |  |  | 0.035 | 0.115 |
| *Micromonas polaris* | RCC2306 | EXP1 | Light-replete | YG-beads | 1 | 2 | 1.7 | 29.0±0.2 |  | 29.0±0.1 | -0.0 |  |  | 0.832 | 0.833 |
| *Micromonas polaris* | RCC2306 | EXP1 | Light-limited | YG-beads | 1 | 2 | 1.7 | 27.2±0.9 |  | 27.7±0.3 | 0.4 |  |  | 0.579 | 0.615 |
| *Micromonas polaris* | RCC2306 | EXP1 | Dark-replete | YG-beads | 2 | 2 | 2.9 | 6.0±0.3 |  | 6.0±0.7 | 0.1 |  |  | 0.883 | 0.891 |
| *Micromonas polaris* | RCC2306 | EXP1 | Dark-limited | YG-beads | 2 | 2 | 2.9 | 5.9±0.3 |  | 5.5±0.2 | -0.3 |  |  | 0.296 | 0.323 |
| *Micromonas polaris* | RCC2306 | EXP1 | Light-replete | YG-beads | 2 | 2 | 0.5 | 45.6±1.1 |  | 46.5±0.1 | 0.9 |  |  | 0.377 | 0.458 |
| *Micromonas polaris* | RCC2306 | EXP1 | Light-limited | YG-beads | 2 | 2 | 0.8 | 40.7±0.3 |  | 40.5±0.2 | -0.2 |  |  | 0.593 | 0.596 |
| *Micromonas polaris* | RCC4298 | EXP1 | Dark-replete | YG-beads | 1 | 2 | 2.6 | 7.8±0.5 |  | 7.7±0.5 | -0.2 |  |  | 0.740 | 0.740 |
| *Micromonas polaris* | RCC4298 | EXP1 | Dark-limited | YG-beads | 1 | 2 | 2.3 | 7.1±0.2 |  | 6.8±0.1 | -0.3 |  |  | 0.147 | 0.150 |
| *Micromonas polaris* | RCC4298 | EXP1 | Light-replete | YG-beads | 1 | 2 | 1.9 | 41.4±0.8 |  | 40.7±0.6 | -0.7 |  |  | 0.429 | 0.433 |
| *Micromonas polaris* | RCC4298 | EXP1 | Light-limited | YG-beads | 1 | 2 | 2.0 | 30.6±0.3 |  | 30.5±0.1 | -0.1 |  |  | 0.598 | 0.618 |
| *Micromonas polaris* | RCC4298 | EXP1 | Dark-replete | YG-beads | 2 | 2 | 0.5 | 3.0±0.1 |  | 3.0±0.6 | -0.0 |  |  | 0.928 | 0.935 |
| *Micromonas polaris* | RCC4298 | EXP1 | Dark-limited | YG-beads | 2 | 2 | 0.5 | 2.0±0.2 |  | 2.3±0.2 | 0.3 |  |  | 0.293 | 0.302 |
| *Micromonas polaris* | RCC4298 | EXP1 | Light-replete | YG-beads | 2 | 2 | 0.2 | 20.4±1.2 |  | 19.5±0.6 | -0.9 |  |  | 0.452 | 0.482 |
| *Micromonas polaris* | RCC4298 | EXP1 | Light-limited | YG-beads | 2 | 2 | 0.2 | 16.1±1.2 |  | 14.9±0.7 | -1.2 |  |  | 0.340 | 0.365 |
| *Micromonas polaris* | RCC2258 | EXP2 | Dark-replete | YG-beads | 1 | 3 | 2.5 | 5.0±0.1 | 4.7±0.1 | 5.1±0.5 | 0.1 | 0.046 | 0.053 | 0.826 | 0.834 |
| *Micromonas polaris* | RCC2258 | EXP2 | Dark-limited | YG-beads | 1 | 3 | 2.2 | 4.4±0.1 | 4.5±0.1 | 4.3±0.1 | -0.1 | 0.772 | 0.772 | 0.339 | 0.340 |
| *Micromonas polaris* | RCC2258 | EXP2 | Light-replete | YG-beads | 1 | 3 | 1.6 | 33.6±0.2 | 33.5±0.1 | 33.2±0.1 | -0.4 | 0.691 | 0.697 | 0.030 | 0.068 |
| *Micromonas polaris* | RCC2258 | EXP2 | Light-replete-AntiB | YG-beads | 1 | 3 | 1.5 | 33.7±0.3 | 34.2±0.3 | 33.8±0.6 | 0.1 | 0.112 | 0.113 | 0.881 | 0.883 |
| *Micromonas polaris* | RCC2258 | EXP2 | Light-limited | YG-beads | 1 | 3 | 1.8 | 33.8±0.8 | 33.6±0.2 | 33.1±0.1 | -0.7 | 0.673 | 0.689 | 0.208 | 0.270 |
| *Micromonas polaris* | RCC2306 | EXP2 | Dark-replete | YG-beads | 1 | 3 | 2.1 | 4.6±0.2 | 4.9±0.2 | 4.6±0.2 | 0.0 | 0.113 | 0.117 | 0.741 | 0.742 |
| *Micromonas polaris* | RCC2306 | EXP2 | Dark-limited | YG-beads | 1 | 3 | 1.9 | 4.4±0.2 | 3.9±0.4 | 4.7±0.3 | 0.3 | 0.172 | 0.190 | 0.279 | 0.290 |
| *Micromonas polaris* | RCC2306 | EXP2 | Light-replete | YG-beads | 1 | 3 | 1.6 | 27.8±1.1 | 28.8±0.4 | 29.2±0.3 | 1.4 | 0.213 | 0.253 | 0.099 | 0.151 |
| *Micromonas polaris* | RCC2306 | EXP2 | Light-replete-AntiB | YG-beads | 1 | 3 | 1.6 | 27.8±0.3 | 29.8±2.5 | 28.7±0.5 | 0.9 | 0.235 | 0.294 | 0.068 | 0.086 |
| *Micromonas polaris* | RCC2306 | EXP2 | Light-limited | YG-beads | 1 | 3 | 1.6 | 28.0±0.2 | 28.0±0.1 | 27.9±0.1 | -0.1 | 0.782 | 0.785 | 0.551 | 0.557 |
| *Micromonas polaris* | RCC4298 | EXP3 | Light-replete | FLBs | 1 | 3 | 1.8 | 24.5±2.3 |  | 24.6±2.7 | 0.0 |  |  | 0.984 | 0.984 |
| *Micromonas polaris* | RCC4298 | EXP3 | Light-replete | YG-beads | 1 | 3 | 1.6 | 35.6±2.2 |  | 34.9±2.0 | -0.7 |  |  | 0.709 | 0.709 |
| *Micromonas polaris* | RCC2306 | EXP5 | Dark-replete | YG-beads | 1 | 2 | 1.1 | 4.4±0.3 |  |  |  |  |  |  |  |
| *Micromonas polaris* | RCC2306 | EXP5 | Light-replete | YG-beads | 1 | 2 | 0.9 | 4.6±0.4 |  |  |  |  |  |  |  |
| *Micromonas polaris* | RCC2306 | EXP5 | Light-replete | YG-beads | 1 | 2 | 0.7 | 4.6±0.0 |  |  |  |  |  |  |  |
| *Micromonas polaris* | RCC2306 | EXP5 | Light-replete | YG-beads | 1 | 2 | 0.1 | 0.4±0.1 |  |  |  |  |  |  |  |
| *Micromonas polaris* | RCC2306 | EXP5 | Light-replete | YG-beads | 1 | 2 | 0.0 | 0.1±0.0 |  |  |  |  |  |  |  |
| *Micromonas polaris* | RCC2306 | EXP5 | Light-replete | YG-beads | 1 | 2 | 4.0 | 6.3±0.5 |  |  |  |  |  |  |  |
| *Micromonas polaris* | RCC2306 | EXP5 | Light-replete | YG-beads | 1 | 2 | 0.6 | 0.8±0.1 |  |  |  |  |  |  |  |
| *Micromonas polaris* | RCC2306 | EXP5 | Light-replete | YG-beads | 1 | 2 | 0.1 | 0.2±0.2 |  |  |  |  |  |  |  |
| *Micromonas polaris* | RCC2306 | EXP5 | Light-replete | YG-beads | 1 | 2 | 1.3 | 4.4±0.2 |  |  |  |  |  |  |  |
| *Micromonas polaris* | RCC2306 | EXP5 | Light-limited | YG-beads | 1 | 2 | 2.1 | 4.4±0.1 |  |  |  |  |  |  |  |
| *Micromonas polaris* | RCC4298 | EXP5 | Light-replete | YG-beads | 1 | 2 | 0.5 | 5.6±0.2 |  |  |  |  |  |  |  |
| *Micromonas polaris* | RCC4298 | EXP5 | Light-replete | YG-beads | 1 | 2 | 0.1 | 0.5±0.2 |  |  |  |  |  |  |  |
| *Micromonas polaris* | RCC4298 | EXP5 | Light-replete | YG-beads | 1 | 2 | 0.0 | 0.1±0.0 |  |  |  |  |  |  |  |
| *Micromonas polaris* | RCC4298 | EXP5 | Light-replete | YG-beads | 1 | 2 | 4.8 | 4.8±1.5 |  |  |  |  |  |  |  |
| *Micromonas polaris* | RCC4298 | EXP5 | Light-replete | YG-beads | 1 | 2 | 0.5 | 0.5±0.5 |  |  |  |  |  |  |  |
| *Micromonas polaris* | RCC4298 | EXP5 | Light-replete | YG-beads | 1 | 2 | 0.1 | 0.2±0.2 |  |  |  |  |  |  |  |
| *Micromonas polaris* | RCC4298 | EXP5 | Dark-replete | YG-beads | 1 | 3 | 1.8 | 7.0±0.1 |  |  |  |  |  |  |  |
| *Micromonas polaris* | RCC4298 | EXP5 | Dark-limited | YG-beads | 1 | 3 | 1.9 | 6.4±0.2 |  |  |  |  |  |  |  |
| *Micromonas polaris* | RCC4298 | EXP5 | Light-replete | YG-beads | 1 | 3 | 1.7 | 24.7±0.4 |  |  |  |  |  |  |  |
| *Micromonas polaris* | RCC4298 | EXP5 | Light-replete-AntiB | YG-beads | 1 | 3 | 1.9 | 23.8±0.7 |  |  |  |  |  |  |  |
| *Micromonas polaris* | RCC4298 | EXP5 | Light-limited | YG-beads | 1 | 3 | 1.8 | 22.5±0.2 |  |  |  |  |  |  |  |

**Table S4.** Comparison of feeding on three different YG-bead sizes (0.5, 1, and 2 *μ*m in diameter) for *M. polaris* (EXP7). The percent of cells with preys (Mean±sd) was measured independently for each bead size and is indicated for each time point (T_0_ and T_40_, where the subscript).

| Species | strain | Treatment | Prey | Replication | Prey/cell ratio | $T_0$ | $T_{40}$ | $T_{40}-T_0$ |
| --- | --- | --- | --- | --- | --- | --- | --- | --- |
| *Micromonas polaris* | RCC2306 | Light-limited | YG-beads 0.5 $\mu\text{m}$ | 2 | 1.0 | 4.09±0.32 | 4.08±0.07 | 0.0 |
| *Micromonas polaris* | RCC2306 | Light-limited | YG-beads 1.0 $\mu\text{m}$ | 2 | 0.9 | 3.27±0.14 | 2.30±0.10 | -1.0 |
| *Micromonas polaris* | RCC2306 | Light-limited | YG-beads 2.0 $\mu\text{m}$ | 2 | 0.7 | 3.54±0.03 | 2.74±0.05 | -0.8 |

**Table S5.** Comparison of Lugol’s iodine and glutaraldehyde fixation, and live (no fixation) measurements of the percent of *M. polaris* cells with YG-beads (EXP6). The percent of cells with preys (Mean±sd) is indicated for each time point after the addition of prey (T_0_ and T_40_, where the subscript corresponds to minutes).

| Species | strain | Treatment | Prey | Fixation | Replication | Prey/cell ratio | $T_0$ | $T_{40}$ | $T_{40}-T_0$ |
| --- | --- | --- | --- | --- | --- | --- | --- | --- | --- |
| *Micromonas polaris* | RCC2306 | Light-replete | YG-beads | No (Live) | 2 | 1.6 | 7.57±0.60 | 7.20±0.28 | -0.4 |
| *Micromonas polaris* | RCC2306 | Light-replete | YG-beads | Lugols iodine | 2 | 1.4 | 5.85±0.39 | 5.58±0.18 | -0.3 |
| *Micromonas polaris* | RCC2306 | Light-replete | YG-beads | Glutaraldehyde | 2 | 1.5 | 6.97±0.39 | 6.62±0.01 | -0.3 |

**Table S6.** Lysosensor experiment (EXP9). Last column shows the Mean±sd Lysosensor green.

| Species | strain | treatment | Unstained or Stained | Replication | Green fluorescence |
| --- | --- | --- | --- | --- | --- |
| *Ochromonas triangulata* | RCC21 | Light-limited | Unstained | 2 | 64.0±1.1 |
| *Ochromonas triangulata* | RCC21 | Light-limited | Stained | 2 | 221.5±40.5 |
| *Micromonas polaris* | RCC2306 | Light-limited | Unstained | 2 | 56.7±2.4 |
| *Micromonas polaris* | RCC2306 | Light-limited | Stained | 2 | 67.8±1.9 |

**Figure S1.**
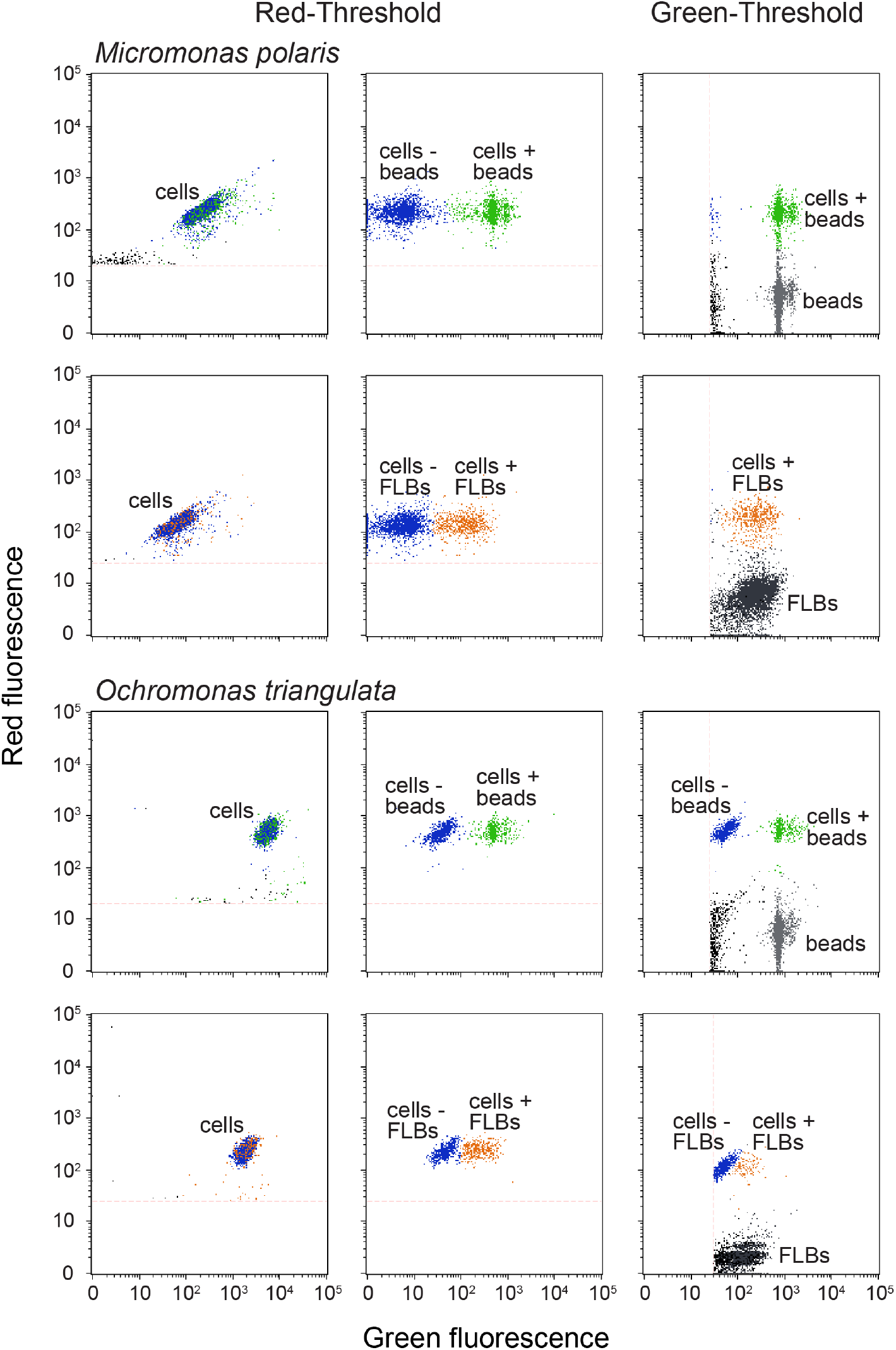
Examples of flow cytograms for *M. polaris* and the positive control *O. triangulata*. Flow cytometry was used to determine the percent of cells with preys (YG-beads and FLBs) in fixed samples (protocol modified from Sherr *et al.* [46]). Data collection was performed with threshold on red (695 ± 50 nm band pass filter) or green fluorescence (525 ± 30 nm band pass filter). Cells that displayed red autofluorescence from chlorophyll as well as green fluorescence were considered to be containing prey (cells with YG-beads in green, cells with FLBs in orange and cells without prey in blue). In addition, to confirm the total concentration of prey added to each experimental flask, the same sample was also run with the threshold on green fluorescence (YG-beads and FLBs in grey and black respectively).

**Figure S2.**
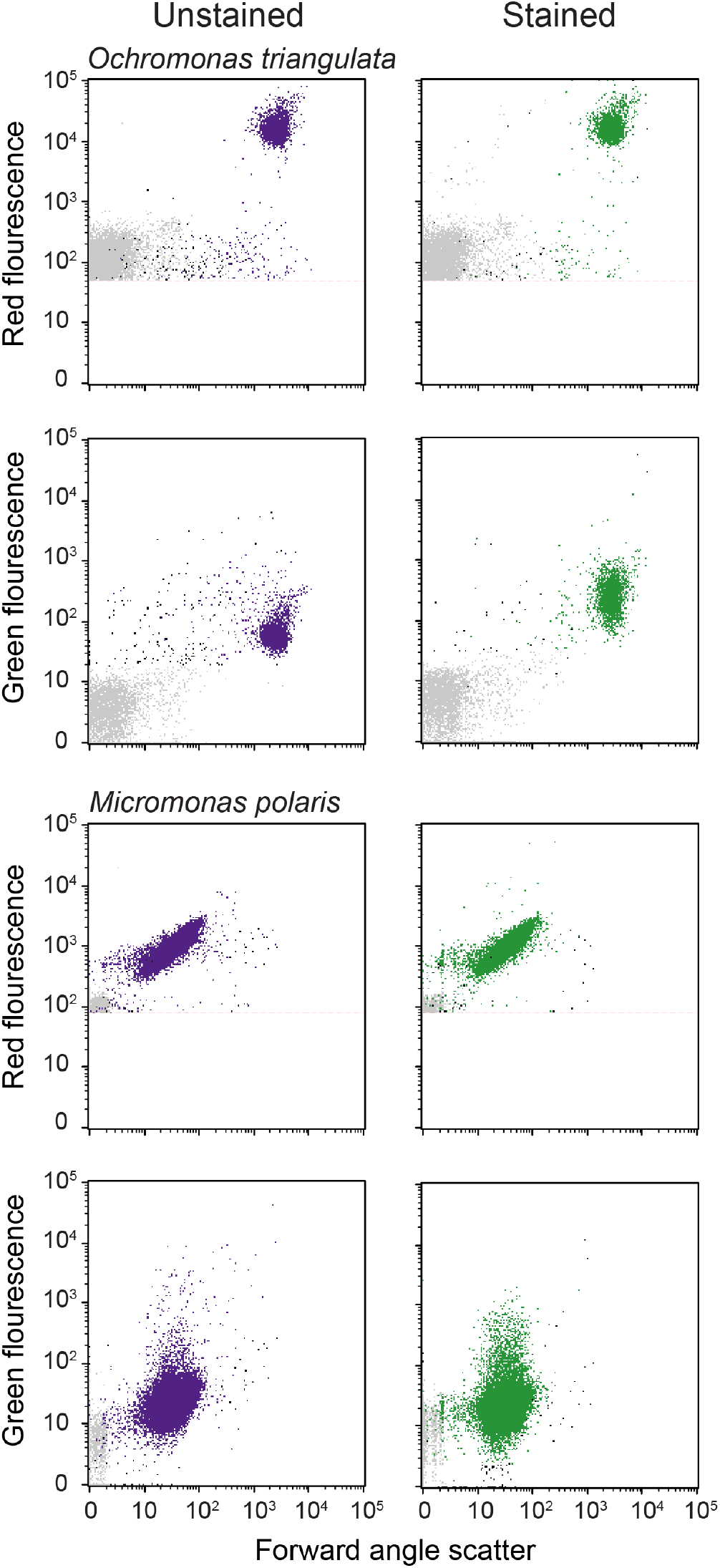
Flow cytograms of *O. triangulata* and *M. polaris* before (purple) and after (green) staining with Lysosensor. Red fluorescence corresponds to chlorophyll fluorescence, while green fluorescence corresponds to autofluorescence before staining or to Lysosensor fluorescence after staining. Green fluorescence clearly increases after Lysosensor staining for *O. triangulata* and not for *M. polaris*.

**Figure S3.**
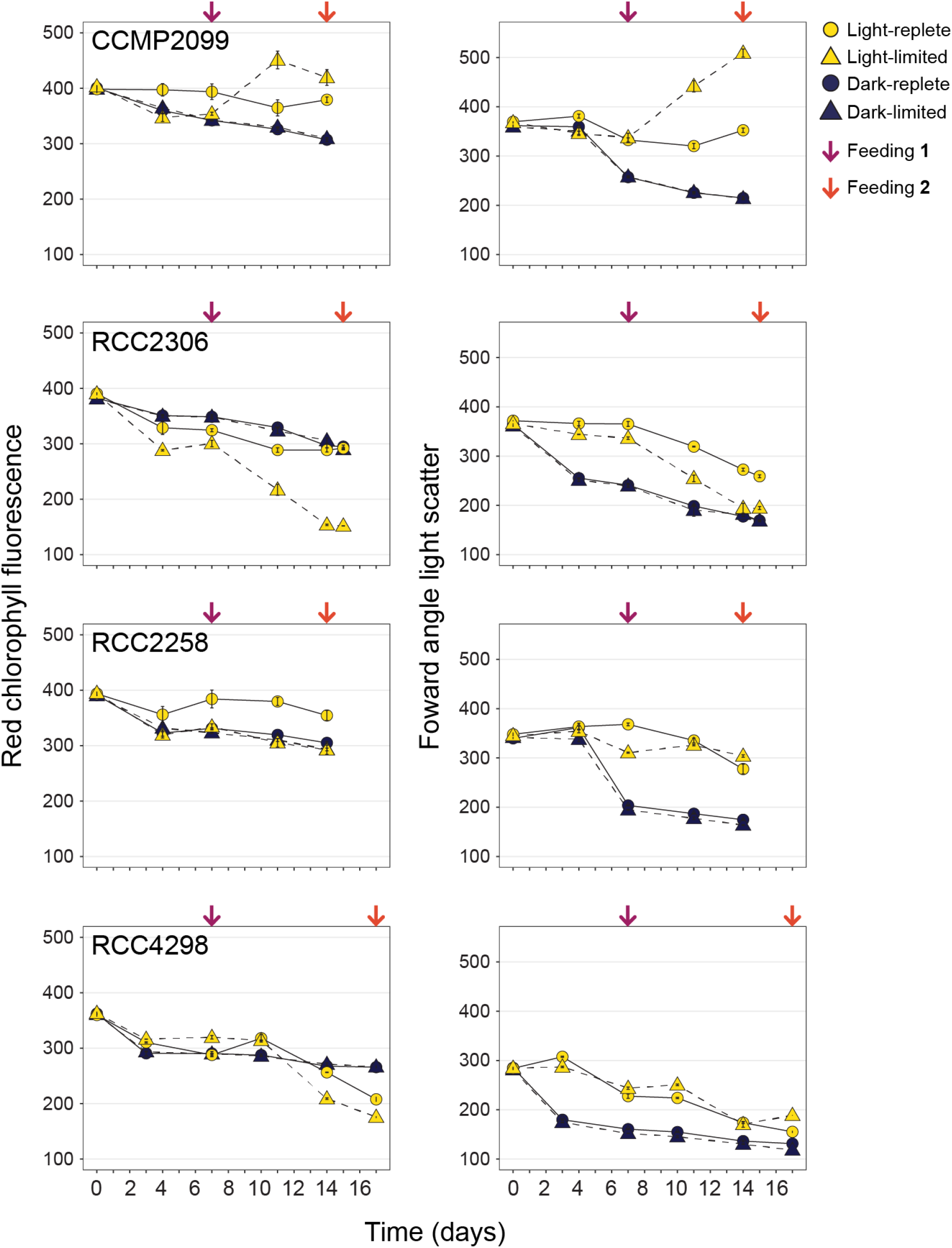
Change in forward scatter and red chlorophyll fluorescence measured by flow cytometry during the experiments reported in Figure 1 (*M. polaris*-EXP1).

**Figure S4.**
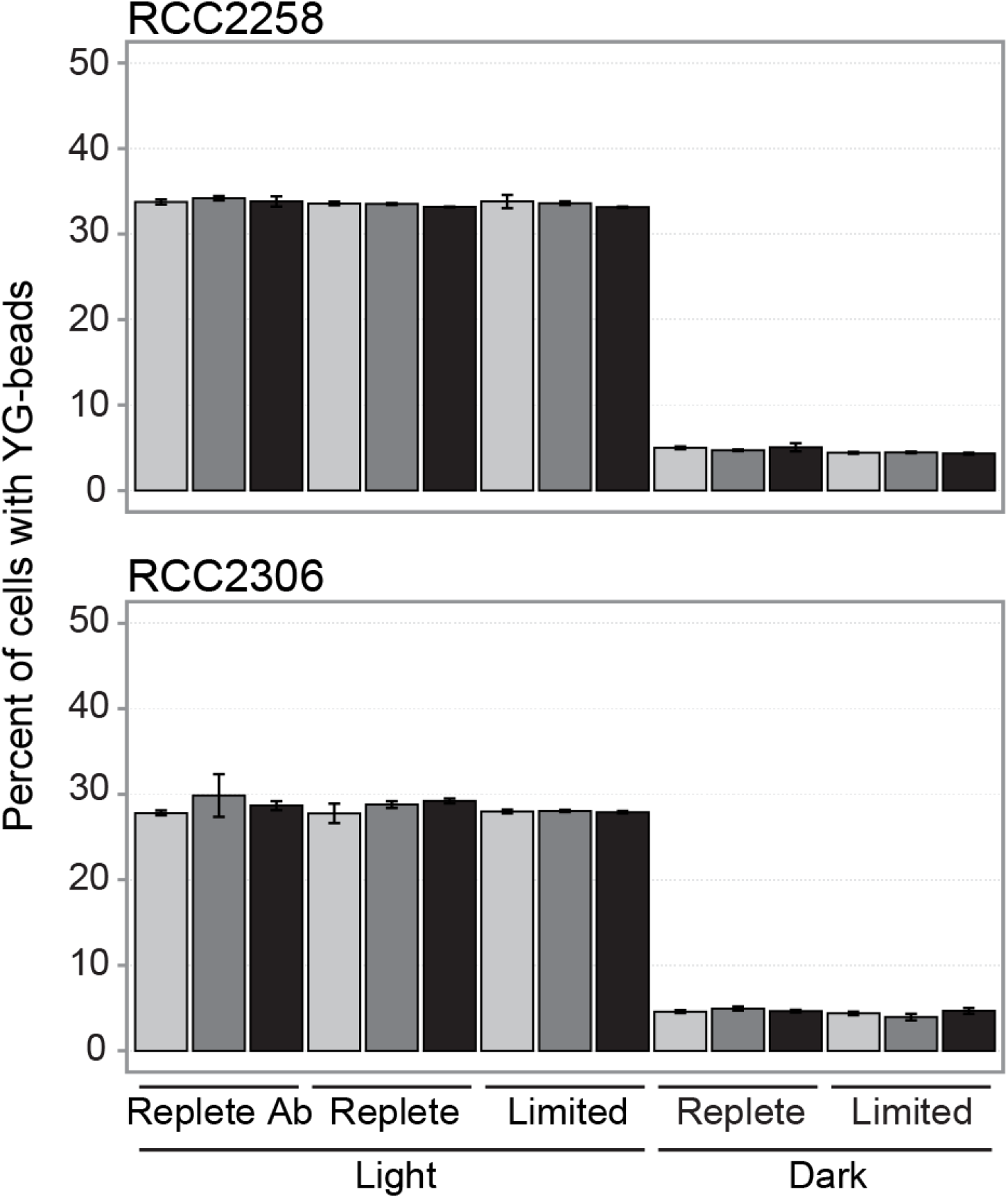
Percent of *M. polaris* cells with YG-beads (*M. polaris*-EXP2) for each strain and treatment. The color of the bars represents the time point (in minutes) after the addition of YG-beads (0 minutes; light grey, 20 minutes; dark grey, 40 minutes; black). Replete Ab correspond to nutrient replete conditions with antibiotics.

**Figure S5.**
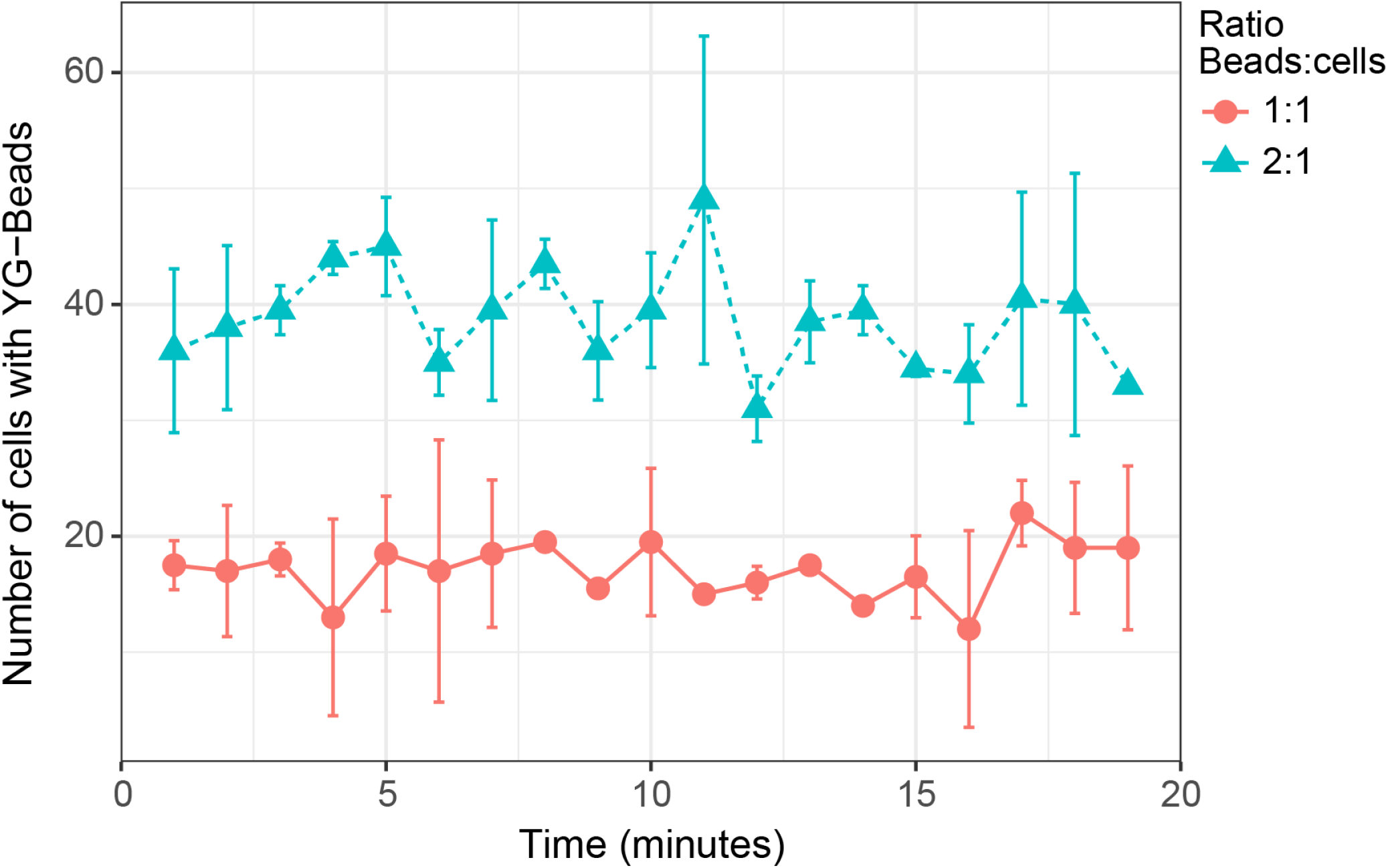
Changes with time in the number of *M. polaris* (strain RCC2306) cells with YG-beads measured by continuously running a live sample for 20 minutes immediately after the addition of YG-beads. Two ratios of beads to cells were tested, 1:1 and 2:1, each in duplicate.

**Figure S6.**
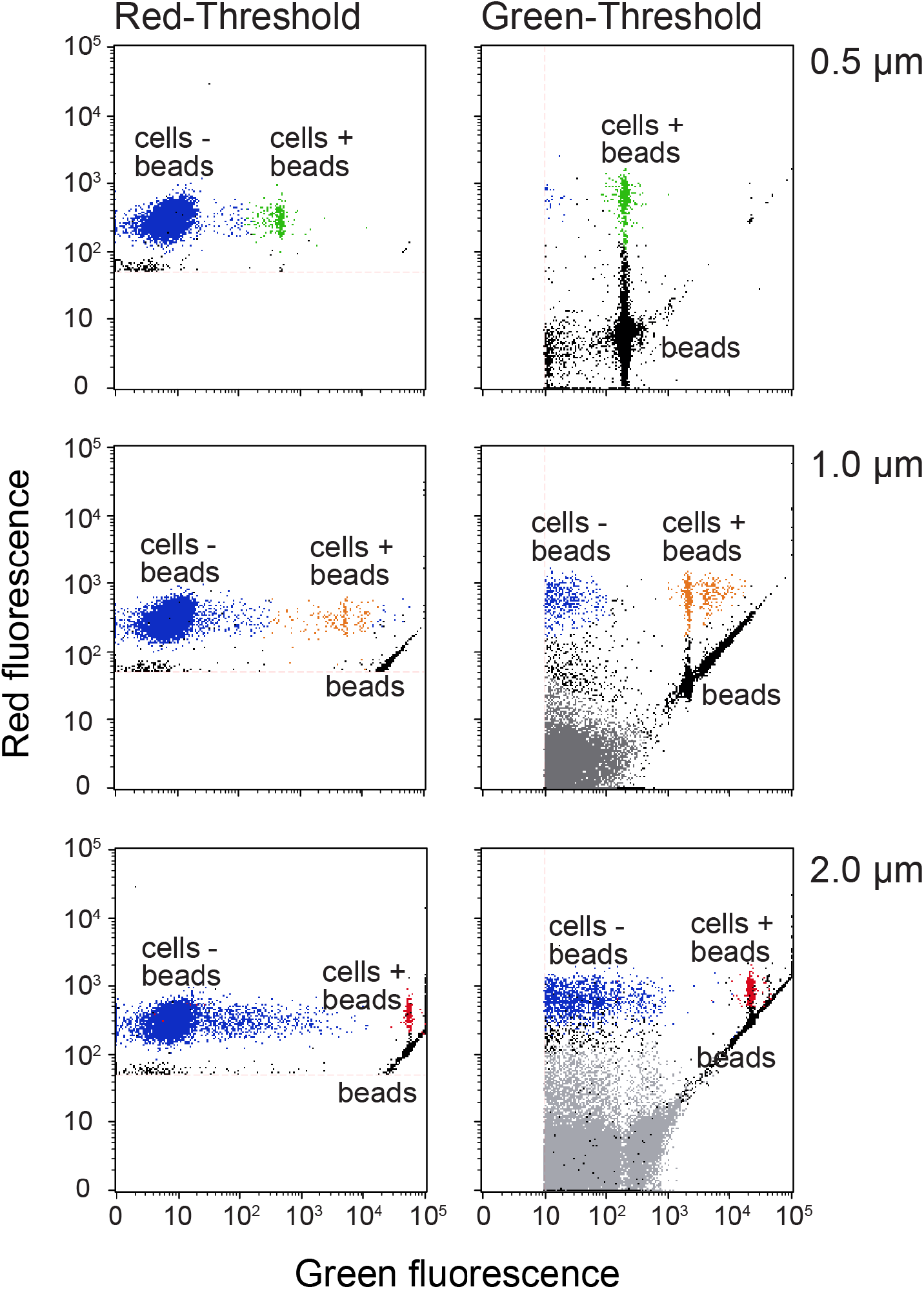
Flow cytograms for *M. polaris* cells incubated with YG-beads of three different sizes (0.5 (green), 1.0 (orange) and 2.0 (red) *μ*m). See legend of Figure S1 for details.

**Figure S7.**
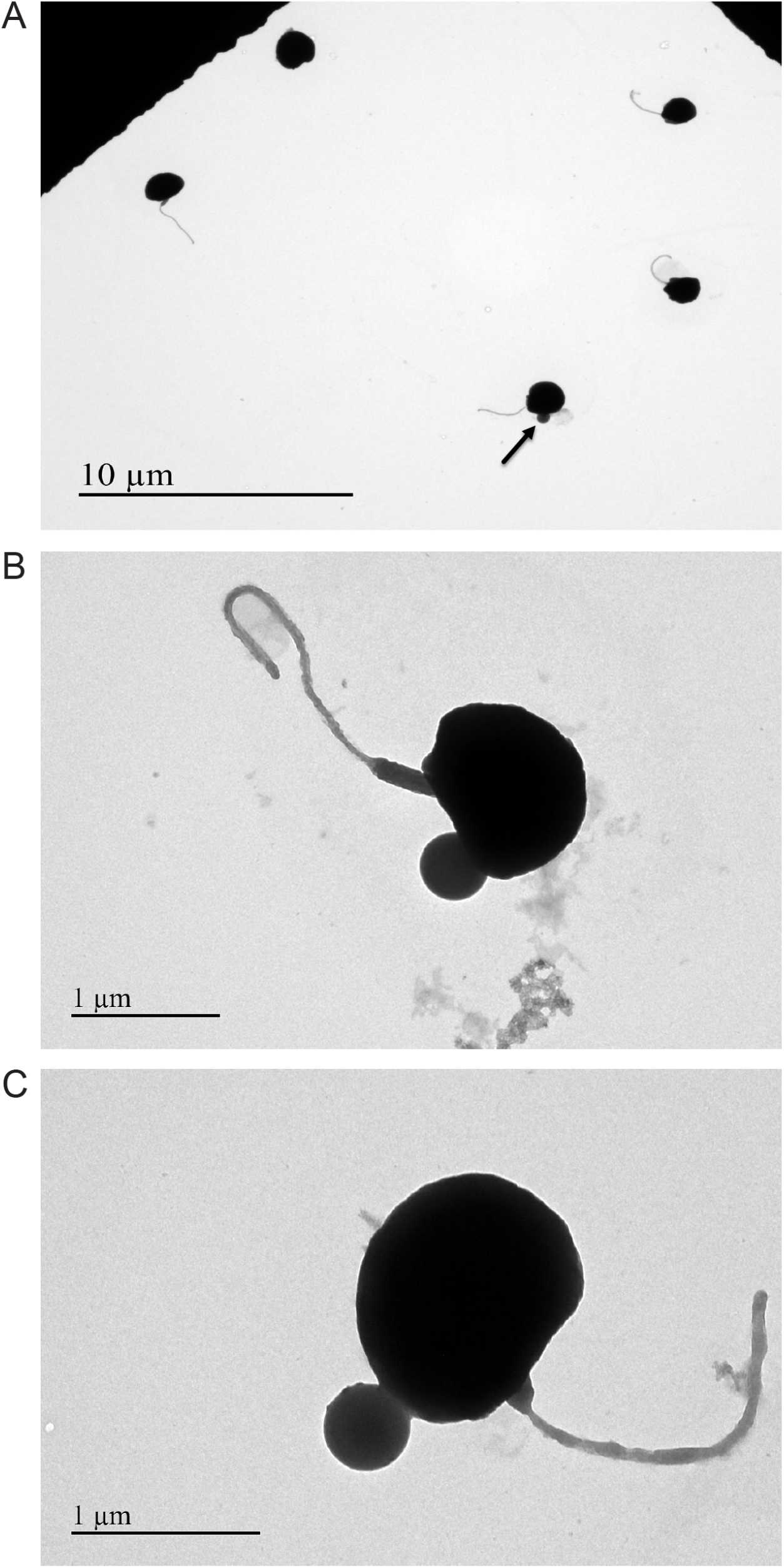
Transmission electron microscopy images of *M. polaris* (strain RCC2306) with YG-beads (0.5 *μ*m) after negative staining. A. Arrow indicates a *M. polaris* cell with a YG-bead. B and C. Close up views of *M. polaris* cells with attached YG-bead.

## Notes

https://github.com/burnsajohn/predictTrophicMode

### Competing Interest Statement

The authors have declared no competing interest.

